# Origin of a novel *CYP20A1* transcript isoform through multiple Alu exaptations creates a potential miRNA sponge

**DOI:** 10.1101/618645

**Authors:** Aniket Bhattacharya, Vineet Jha, Khushboo Singhal, Mahar Fatima, Dayanidhi Singh, Gaura Chaturvedi, Dhwani Dholakia, Rintu Kutum, Rajesh Pandey, Trygve E. Bakken, Pankaj Seth, Beena Pillai, Mitali Mukerji

**Author notes:** Equal contribution (Aniket Bhattacharya, Vineet Jha, Khushboo Singhal).

## Abstract

**Background:** Primate-specific Alus contribute to transcriptional novelties in conserved gene regulatory networks. Alu RNAs are present at elevated levels in stress conditions and consequently leads to transcript isoform specific functional role modulating the physiological outcome. One of the possible mechanisms could be Alu nucleated mRNA-miRNA interplay.

**Result:** Using combination of bioinformatics and experiments, we report a transcript isoform of an orphan gene, *CYP20A1* (*CYP20A1_Alu-LT*) through exaptation of 23 Alus in its 9kb 3’UTR. *CYP20A1_Alu-LT*, confirmed by 3’RACE, is an outlier in length and expressed in multiple cell lines. We demonstrate its presence in single nucleus RNA-seq of ∼16000 human cortical neurons (including rosehip neurons). Its expression is restricted to the higher primates. Most strikingly, miRanda predicts ∼4700 miRNA recognition elements (MREs; with threshold< −25kcal/mol) for ∼1000 miRNAs, which have majorly originated within the 3’UTR-Alus post exaptation. We hypothesized that differential expression of this transcript could modulate mRNA-miRNA networks and tested it in primary human neurons where *CYP20A1_Alu-LT* is downregulated during heat shock response and upregulated upon HIV1-Tat treatment. *CYP20A1_Alu-LT* could possibly function as a miRNA sponge as it exhibits features of a sponge RNA such as cytosolic localization and ≥10 MREs for 140 miRNAs. Small RNA-seq revealed expression of nine miRNAs that can potentially be sponged by *CYP20A1_Alu-LT* in neurons. Additionally, *CYP20A1_Alu-LT* expression was positively correlated (low in heat shock and high in Tat) with 380 differentially expressed genes that contain cognate MREs for these nine miRNAs. This set is enriched in genes involved in neuronal development and hemostasis pathways.

**Conclusion:** We demonstrate a potential role for *CYP20A1_Alu-LT* as miRNA sponge through preferential presence of MREs within Alus in a transcript isoform specific manner. This highlights a novel component of Alu-miRNA mediated transcriptional modulation leading to physiological homeostasis.

## Introduction

Nearly half of the human genome is occupied by transposable elements (TEs) (1). These have been shown to fine-tune conserved gene regulatory networks in a lineage specific manner (2–4). Depending upon the context, they contribute to gene expression divergence through large scale transcriptional rewiring (3,5,6). Primate specific Alu retrotransposons, which occupy ∼11% of the human genome, are major players in this process (1,7). These provide non-canonical transcription factor binding sites and other regulatory sites that govern epigenetic modifications as well as provide cryptic splice sites that lead to alternative splicing or differential mRNA stability (5,8–14). Alu-derived exons exhibit lineage specificity with high transcript inclusion levels and have much higher rates of evolution (15–17).

Nearly 14% of the human transcriptome contain exonized Alus, predominantly in the principal isoforms of genes. Exonization is frequently reported in genes that have arisen *de novo* in primates, with most of the events in 3’UTRs (18,19). Such exonized Alus can increase the regulatory possibilities for a transcript, in a spatio-temporal manner, through antisense, miRNAs, A-to-I RNA editing, alternative splicing and enhancers. We have shown that transcript isoform dynamics i.e., the relative proportion of exonized versus non-exonized isoforms of a gene, could be modulated by these events in viral recovery and stress response (19,20). Besides, Alus provide substrates for other regulatory events such as gain of poly-A sites, AU-rich motifs and miRNA recognition elements (MREs) that can result in alternative polyadenylation, mRNA decay or translation stalling; and formation of specific secondary structures (9,21–25). We have also reported that Alu-MREs can alter transcript isoform dynamics during stress response and some of these sites seem to be evolving in humans (20).

In our earlier study on 3177 Alu-exonized genes, we reported the co-occurrence of *cis* Alu antisense and A-to-I editing at the level of single Alu exons in 319 genes. Enrichment analysis revealed genes related to apoptosis and lysosomal processes to be enriched (19). During mapping of lineage specific events in these genes, we observed a gene, *CYP20A1* that has acquired an unusually long 3’ UTR due to the exaptation of 23 Alus belonging to different subfamilies. This has led to the creation of a novel transcript isoform that we named *CYP20A1_Alu-LT*. Analysis revealed that its 3’UTR contains target sites for ∼1000 miRNAs, predominantly within Alu, with ≥10 MREs for 140 miRNAs. Based on multiple shared features, we hypothesize that *CYP20A1_Alu-LT* could function as miRNA sponge that has originated from repetitive sequences. We demonstrate its regulatory potential in primary neurons wherein the presence of miRNAs and the expression of *CYP20A1_Alu-LT* correlates with the expression of RNA-seq derived genes (in enriched processes) that share cognate MREs. Our results highlight a miRNA sponge, derived from Alus, which could orchestrate systemic changes in associated miRNA regulatory networks.

## Results

### *CYP20A1* contains a unique 3’UTR with Alu-driven divergence

Our previous work had identified 319 Alu exonized genes wherein co-occurrence of regulatory events coalesced within Alus (19). Literature mining revealed that 91 out of these 319 genes map to apoptosis and nearly 75% of them cluster around three discrete hubs: cell cycle-DNA damage response (p53 hub; 31 genes), mitochondrial events (mito hub; 22 genes) and proteostasis (ubi hub; 15 genes) **(Supplementary Information S1 and Table S1)**. As the majority of these exonization events occur in the 3’UTRs of transcripts, which can modulate miRNA regulatory networks, we focused on identifying specific events in the 3’UTRs of these genes.

We found a transcript isoform of *CYP20A1* gene (referred to as *CYP20A1_Alu-LT* hereafter) that has an 8.93kb long 3’UTR, 65% of which is derived from the exonization of 23 Alus **(Figure 1a)**. Since this transcript has an unusual density of Alus across the length of the UTR, we characterized it further for regulatory potential. Amongst the eight transcripts of human *CYP20A1* annotated in NCBI, experimental evidence is available only for *CYP20A1_Alu-LT* (NM_177538), the longest isoform (10.94 kb). It is an outlier in terms of its 3’UTR length as it occupies the 85th position in the genome-wide length distribution of 3’UTRs **(Figure 1b)**. Such extended 3’UTRs are extremely rare and even among Alu-exonized genes, less than 3% have UTRs longer than 6kb **(Supplementary Figure S1)**. The length and enrichment do not seem to correlate with the density of exonized Alus (r = 0.25) compared to the genomic average of 5.42 exonization events/ 3’UTR. The Alus in this transcript belong to subfamilies of different evolutionary ages, suggesting that their insertion could have happened over a period.

**Figure 1:**
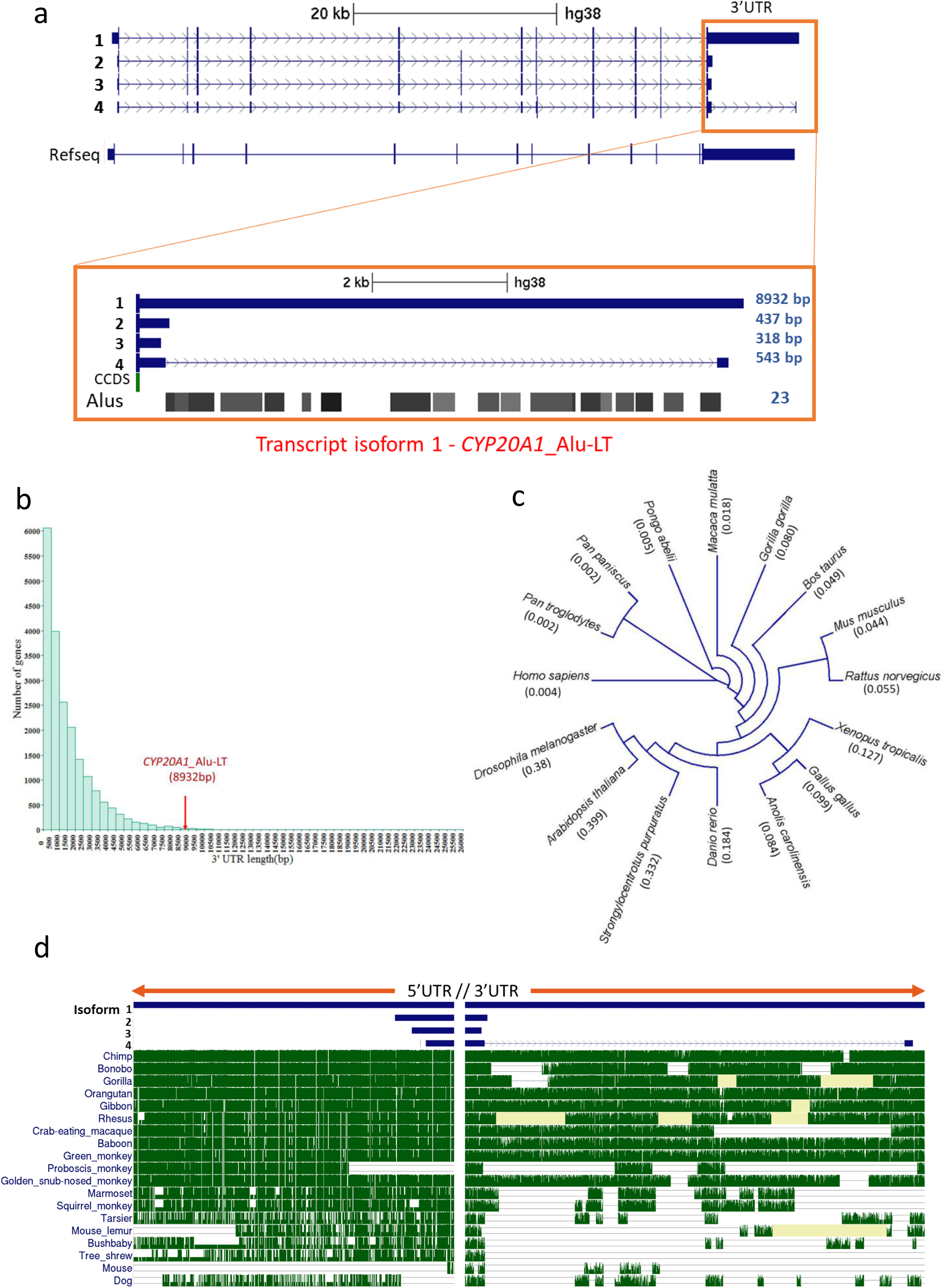
*CYP20A1* contains a unique 3’UTR with Alu-driven divergence. (a) UCSC tracks representing the four transcript isoforms of *CYP20A1* with varying 3’UTR length. Only isoform 1 (NM_177538) contains the full length 8932bp 3’UTR (*CYP20A1_Alu-LT*). The RepeatMasker track shows this 3’UTR harbours 23 Alu repeats from different subfamilies. (b) Genome-wide analysis of length distribution of 3’UTR reveals *CYP20A1_Alu-LT* to be an outlier. Mean and median 3’UTR lengths were 1553bp and 1007bp, respectively. (c) Cladogram of CYP20A1 protein sequence divergence among different classes of vertebrates. At the protein level this gene seems to have diverged minimally. (d) DNA level conservation analysis of 5’UTR and 3’UTR among 20 mammals reveals that 5’UTR is well conserved among all primate lineages, suggesting that divergence is unique to 3’UTR (also see **Supplementary Figure S3**).

### Genomic region proximal to *CYP20A1_Alu-LT* 3’UTR is relatively well conserved

The coding region of *CYP20A1_Alu-LT* is remarkably well conserved among vertebrates, both at the sequence level as well as the length of the mature protein **(Table 1)**. The chimpanzee, macaque and mouse *CYP20A1* code for the same 462-470aa protein as in humans although their annotated transcript orthologs range between 1-3kb. Multiple sequence alignment across vertebrates reveals a strong conservation at both the N and the C terminals **(Supplementary Figure S2)**; of the first 100aa, 62 are completely conserved while 18 contain lineage specific substitutions with residues that have similar functional groups. This is also corroborated by the minimal evolutionary divergence across vertebrate CYP20A1 proteins **(Figure 1c)** and a strong purifying selection in CDS (Ka/Ks ∼0.2 in mammals and <0.1 in non-mammalian vertebrates) **(Table 1)**.

**Table 1:**
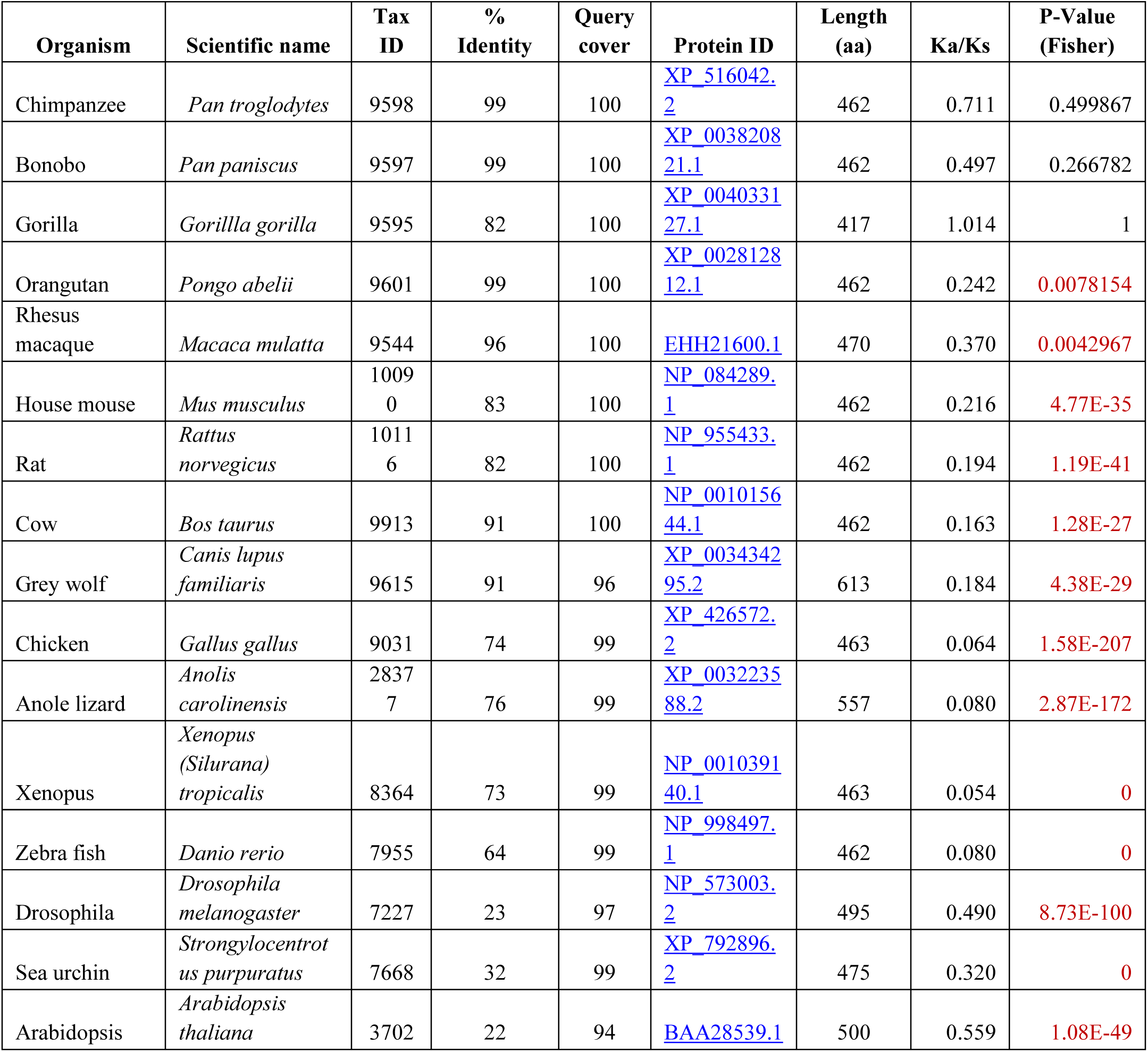
CYP20A1 protein sequence conservation across vertebrates. CYP20A1 protein sequences among different vertebrate classes were compared using NCBI pBLAST. Drosophila, sea urchin and Arabidopsis were used as the evolutionary outgroups. All the pairwise comparisons are done with respect to the 462aa human CYP20A1 protein (NP_803882) and a query cover of ∼95% was obtained in each case. CYP20A1 is well conserved among vertebrates. Except for the three great apes (chimpanzee, bonobo and gorilla), all the Ka/Ks comparisons are significant (Fisher’s exact test; p<0.01; marked in red). Lesser the value of Ka/Ks, the more stringent is the negative selection operative on the protein i.e., fewer non-synonymous substitutions are tolerated in it.

| Organism | Scientific name | Tax ID | % Identity | Query cover | Protein ID | Length (aa) | Ka/Ks | P-Value (Fisher) |
| --- | --- | --- | --- | --- | --- | --- | --- | --- |
| Chimpanzee | *Pan troglodytes* | 9598 | 99 | 100 | XP_516042.2 | 462 | 0.711 | 0.499867 |
| Bonobo | *Pan paniscus* | 9597 | 99 | 100 | XP_003820821.1 | 462 | 0.497 | 0.266782 |
| Gorilla | *Gorilla gorilla* | 9595 | 82 | 100 | XP_004033127.1 | 417 | 1.014 | 1 |
| Orangutan | *Pongo abelii* | 9601 | 99 | 100 | XP_002812812.1 | 462 | 0.242 | 0.0078154 |
| Rhesus macaque | *Macaca mulatta* | 9544 | 96 | 100 | EHH21600.1 | 470 | 0.370 | 0.0042967 |
| House mouse | *Mus musculus* | 10090 | 83 | 100 | NP_084289.1 | 462 | 0.216 | 4.77E-35 |
| Rat | *Rattus norvegicus* | 10116 | 82 | 100 | NP_955433.1 | 462 | 0.194 | 1.19E-41 |
| Cow | *Bos taurus* | 9913 | 91 | 100 | NP_001015644.1 | 462 | 0.163 | 1.28E-27 |
| Grey wolf | *Canis lupus familiaris* | 9615 | 91 | 96 | XP_003434295.2 | 613 | 0.184 | 4.38E-29 |
| Chicken | *Gallus gallus* | 9031 | 74 | 99 | XP_426572.2 | 463 | 0.064 | 1.58E-207 |
| Anole lizard | *Anolis carolinensis* | 28377 | 76 | 99 | XP_003223588.2 | 557 | 0.080 | 2.87E-172 |
| Xenopus | *Xenopus (Silurana) tropicalis* | 8364 | 73 | 99 | NP_001039140.1 | 463 | 0.054 | 0 |
| Zebra fish | *Danio rerio* | 7955 | 64 | 99 | NP_998497.1 | 462 | 0.080 | 0 |
| Drosophila | *Drosophila melanogaster* | 7227 | 23 | 97 | NP_573003.2 | 495 | 0.490 | 8.73E-100 |
| Sea urchin | *Strongylocentrotus purpuratus* | 7668 | 32 | 99 | XP_792896.2 | 475 | 0.320 | 0 |
| Arabidopsis | *Arabidopsis thaliana* | 3702 | 22 | 94 | BAA28539.1 | 500 | 0.559 | 1.08E-49 |

On the other hand, 3’UTR extension in *CYP20A1_Alu-LT* seems to be mediated by the primate specific insertion of Alus. Its orthologs in mouse, rat and zebrafish are extremely short (within 1kb). In mouse, we observe a sparse presence of two B1 SINEs, one each of simple repeat and low complexity repeat whereas the zebrafish 3’UTR lacks repeats altogether. The longest annotated *CYP20A1* transcripts for mouse (NM_030013.3), rat (NM_199401.1) and zebrafish (NM_213332.2) are 2.27, 2.03 and 1.79kb, respectively. The 5’UTR appears to be well conserved across the primate lineage (except lemur and proboscis monkey); however, the divergence in the 3’UTR, as evident from Jukes Cantor measure, increases as we move from the great apes to rhesus macaque and is primarily contributed by the Alus, with the breakpoints mostly coinciding with an Alu insertion **(Figure 1d)**. It shows maximum divergence from mouse that was treated as a non-primate evolutionary out-group. To control for the length difference between the 5’ and 3’UTRs, we also checked for conservation in the 10kb region upstream of the first transcription start site (TSS) of *CYP20A1* and 10kb downstream of transcription end site (TES) and found it to be almost perfectly conserved among the higher primates, except for some New World monkeys **(Supplementary Figure S3)**. Taken together, these observations suggest that insertion of exonized Alus might have contributed to the specific divergence of this 3’UTR, in a genomic region that is otherwise conserved, at least among the higher primates.

Since this UTR seems to have appeared relatively late in the primate evolution, we tested if it carries variations that can differentiate modern human populations. Among the 23 SNPs in *CYP20A1* 3’UTR (16 within Alus), 11 have average heterozygosity scores >0.2, some of them as high as 0.48. We analyzed the data from 1000 Genomes Phase I and found significant differentiation for seven of these SNPs with global F_ST_ values ranging between 0.2-0.4 **(Supplementary Table S2)**. Interestingly, we also found a GWAS SNP (rs11888559, C/T, T=0.237/1187) in this UTR, which is associated with height in Filipino women (1) with a global F_ST_ of 0.36, rs11888559 differentiates the east Asian (CHB) and European (CEU) from the ancient African (YRI) population. Expectedly, it also exhibits high-derived allele frequencies (DAFs) in these populations (0.81 and 0.95 in CHB and CEU, respectively). We also found another SNP rs7577078, within Alu, with high DAFs in all the three populations.

### Characterization of *CYP20A1_Alu-LT*

We next investigated whether the full-length transcript containing this uniquely diverged 3’UTR is actually transcribed. As 2/3^rd^ of this 3’UTR comprise repetitive sequences, it was a challenge to capture the full-length transcript in expression arrays or map it uniquely from sequencing reads. Moreover, there are differences in annotations regarding the full-length 3’UTR-containing isoform in various genomic portals **(Supplementary information, S1)**. Therefore, we designed eleven pairs of primers spanning the entire length of the transcript and experimentally confirmed the expression of *CYP20A1_Alu-LT*. We validated three of its amplicons by Sanger sequencing to negate spurious amplification from other Alu-rich loci in the genome (**Figure 2a**, also see Supplementary information, S1 for detailed method).

**Figure 2:**
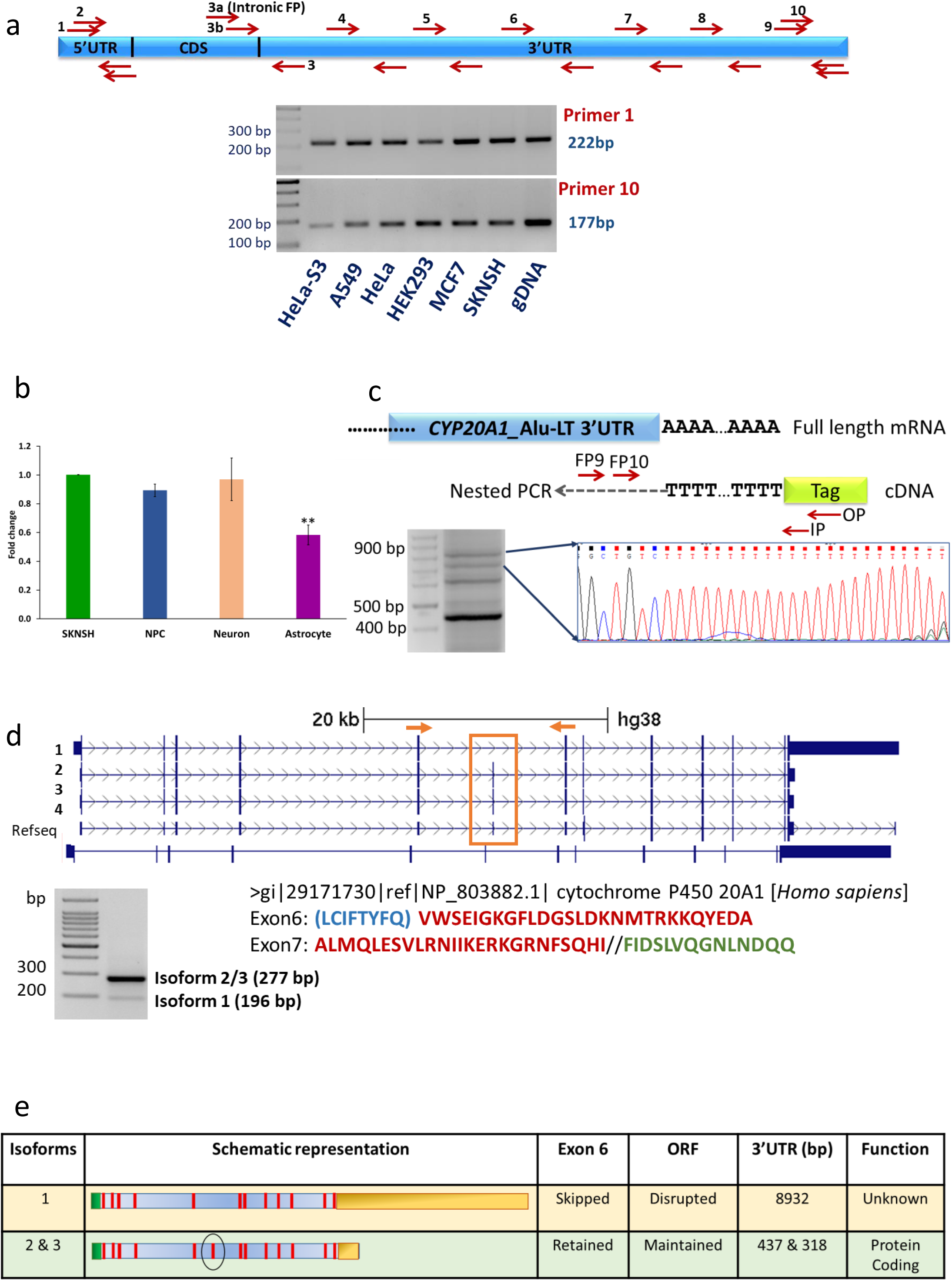
*CYP20A1_Alu-LT* is expressed and may be a long non-coding RNA. (a) A schematic representation of the primers designed on the *CYP20A1_Alu-LT* to encompass 5’UTR and full length 3’UTR. To check for full length expression of transcript, cDNA from multiple cell lines of different tissue origin was used for amplification. Representative gel images of this isoform expression, via amplification from starting of 5’UTR and end of 3’UTR is shown by primer pairs 1 and 10, respectively. Amplicons (1, 5 and 10) were also confirmed by Sanger sequencing. * genomic DNA was used as positive control for PCR. (b) RT-qPCR for *CYP20A1_Alu-LT* expression in cancerous and non cancerous cell types of neuronal origin. Fold change was calculated with respect to SK-N-SH, after normalization with the geometric mean of expression values from *β-actin, GAPDH* and *18S rRNA*. The error bars represent the SD of three biological replicates and the average of three technical replicates were taken for each biological replicate (** → p<0.01; Student’s t test). (c) 3’RACE confirms the expression of the full length transcript. The schematic depicts the oligo(dT) (attached to a tag sequence) primed reverse transcription, followed by nested PCR. The amplification products corresponding to the bands below 900bp and above 700bp mapped to *CYP20A1_Alu-LT* 3’UTR, suggesting that the full length transcript is expressed in untreated MCF-7 cells (n=3). (d) Differentiating the *CYP20A1_Alu-LTR* transcript from other isoforms. The schematic highlights the skipped exon 6 and the position of flanking primers on shared exons. The presence of at least two different types of transcripts was confirmed. 277bp amplicon corresponds to isoform(s) that contain exon 6, but have shorter 3’UTRs (isoforms 2 and 3 in **figure 2e**) and 196bp amplicon corresponds to the long-3’UTR isoform (isoform 1). None of the six translation frames of the long 3’UTR isoform match with the annotated protein. The amino acids marked in red are common to both isoform 2 and 3, blue exclusive to isoform 3 and green represents the sequence from isoform 1. (e) Schematic representation summarizing the differences between *CYP20A1* transcript isoform 1 (*CYP20A1_Alu-LT*) and isoforms 2 and 3.

We observed variable expression of *CYP20A1_Alu-LT* in the six cell lines that we had initially tested **(Figure 2a)**. Since we had used cancerous cell lines, its expression could potentially be attributed to the aberrant transcriptional profiles in cancer (26). To delineate if *CYP20A1_Alu-LT* expression is due to the cancerous state of the cells, we compared its expression in a neuroblastoma cell line (SK-N-SH) with those in primary neuron, glia (astrocyte) and neural progenitor cells (NPC). Neuroblastoma shares features with both mature neurons and NPCs, but is distinct from glia and we found that *CYP20A1_Alu-LT* expression differs significantly only between glia and SK-N-SH **(Figure 2b)** but not in neurons or NPCs. These suggest that our observations in the cancerous cell lines are unlikely to be artifactual.

We selected MCF-7, a breast adenocarcinoma cell line, for some of our subsequent experiments as it has been extensively used for drug screening and studying the effect of xenobiotics on different CYP family genes (27,28). The copy number of *CYP20A1* is not altered in this line (2n=2) (29). We performed 3’RACE to determine the exact transcription termination site for *CYP20A1_Alu-LT*. This was followed by nested PCR and amplicon sequencing to confirm *CYP20A1_Alu-LT* as a bonafide RNA transcript **(Figure 2c, Supplementary information, S3)**. Our findings are further supported by the TargetScan (release 7.2) which builds on the longest Gencode 3’UTR and reports even longer, 12.85kb UTR (ENST000000356079.4). The algorithm calculates 3’UTR length based on 3P-seq tags, accounting for the usage of mRNA cleavage and splice sites, normalized across multiple tissues.

### Exon skipping differentiates *CYP20A1_Alu-LT* from the protein coding isoforms

We observed that the expression of *CYP20A1_Alu-LT* is rather low although CYP20A1 protein is relatively abundant **(Supplementary Figure S4)**, suggesting that other isoforms may contribute to protein levels. When we compare *CYP20A1_Alu-LT* with the shorter 3’UTR containing isoforms, we observe a skipping of the sixth exon in this transcript. Using primers encompassing the sixth exon, we could distinguish between the transcripts; with the larger isoform of 196bp amplicon and shorter ones of 277bp which also show a much higher expression **(Figure 2d)**.

In order to assess the relative contribution of different isoforms to the overall expression of *CYP20A1*, we used RNA-seq data from 15928 single nuclei derived from the different layers of human cerebral cortex (30). NM_177538 (*CYP20A1_Alu-LT*) is expressed in 75% of the nuclei whereas all the other RefSeq isoforms are found in <1% (cut-off CPM≥50). There are 7038, 5134 and 1841 single nuclei in which NM_177538 (but no other isoform) is expressed with ≥10, 50 and 100 reads, respectively **(Supplementary Table S3)**. Interestingly, it is expressed in rosehip neurons - a highly specialized cell type in humans, suggestive of its functional relevance **(Supplementary Table S4)** (31).

Although the long 3’UTR transcript is annotated as the principal isoform, its expression level did not correlate with CYP20A1 protein that is relatively abundant in MCF-7 cells **(Supplementary Figure S4)**. To probe further, we performed *in silico* translation of all *CYP20A1* isoforms in six-frames and compared them to the annotated human CYP20A1 protein. The two short 3’UTR isoforms matched – one perfectly and another with an additional amino acid stretch **(Figure 2d)**, but the *CYP20A1_Alu-LT* goes out of frame in the sixth and seventh exon and BLAST analysis of the human proteome does not report any hits with the truncated 24 amino acid peptide. Taken together, these data suggest that the *CYP20A1_Alu-LT* is unlikely to be coding for CYP20A1 protein and may represent a novel non-coding transcript isoform originating from the same locus. This may be a case of evolutionary sub-functionalization of a gene into two different classes of transcripts that might have evolved for different functions **(Figure 2e)** (30).

### *CYP20A1_Alu-LT* expression in non-human primates

Among the non-human primates, we did not find any annotated transcripts beyond 3kb from this locus. Our preliminary analyses of expression data, from public databases, of chimpanzee and macaque (prefrontal cortex, CD4+ T cells) did not yield any reads mapping to this 9kb 3’UTR. Subsequently, we checked for *CYP20A1-Alu-LT* expression in the reference transcriptomes of non-human primates (http://www.nhprtr.org) (31). Total RNA reads derived from 157 libraries of 14 non-human primate species show consistent mapping pattern on *CYP20A1* 3’UTR. Mapping is higher in the neighboring coding exons, but the pattern is consistent across different tissues and the number of reads comparable, with a slightly higher expression in kidney and lungs. In chimpanzee, reads are evenly distributed across the length of the entire 3’UTR; however, distribution is patchy in the other Old world monkeys (with peaks mostly in the non-repeat regions). Expression is minimal in New world monkeys (marmoset, squirrel monkey) and completely absent in lemur, although the adjoining coding exons show comparable expression, suggesting that *CYP20A1_Alu-LT* is expressed only in the higher primates **(Supplementary Information S3)**.

### *CYP20A1_Alu-LT* 3’UTR as an evolving miRNA regulatory hub

Our earlier study had revealed that 3’UTR exonized Alus could provide novel miRNA binding sites thereby making the transcripts amenable to miRNA mediated post-transcriptional regulation (20). We explored whether this recently diverged, Alu-rich 3’UTR could have evolved as a regulatory hub. We first checked whether the 3’UTR of *CYP20A1_Alu-LT* is also targeted by miRNAs. A query in miRTarBase (release 6.0) (32) revealed that *CYP20A1_Alu-LT* 3’UTR had predicted target sites for 169 miRNAs (supported by microarray/ sequencing data), of which 46 were listed as *functional miRNAs* in FuncMir (miRDB) **(Supplementary Table S5)**. Most strikingly, ∼50% of these are either primate-specific or human-specific miRNAs (microRNAviewer) (33). The occurrence of target sites for human-specific miRNAs in this recently evolved UTR prompted us to carry out further in-depth analysis of miRNA recognition elements (MREs).

Since our 3’UTR has diverged across the vertebrate phylogeny, we did not consider algorithms that employ evolutionary conservation as prediction criteria. Many algorithms which predict target sites based on seed sequence matches also seem to have limitations as the length and position of the seed sequence is variable amongst miRNAs (34,35). In order to reduce false MRE prediction in non-conserved regions, we used miRanda that employs a two-step strategy: sequence complementarity, followed by thermodynamic stability of the predicted miRNA-mRNA duplex (36). Using stringent cut-off criteria, we obtained a total of 4742 MREs for 994 miRNAs, 4500 of which overlap with Alus (4382 MREs, if a conservative estimate of >50% overlap is considered) **(Supplementary Table S6)**.

These 4742 MREs span the entire length of the 3’UTR along with several high density pockets in Alu regions **(Figure 3a)**. The 23 exonized Alus mainly belong to Alu S and J family and are from 13 different subfamilies - AluSx, AluSp, AluSc, AluSz6, AluSq2, AluSx3, AluSc8, AluSx1, AluSz, AluSg, AluJo, AluJb and AluJr. Their sequence of insertion into the 3’UTR of *CYP20A1_Alu-LT* is represented in figure 3a in 5’ to 3’ direction from top to bottom of the circos plot. The 994 miRNAs were grouped on the basis of numbers of MREs present in the 3’UTR of *CYP20A1_Alu-LT*. MREs are grouped in range of 1-5, 6-10, 11-20, 21-43, for group 1 (G1), group 2 (G2), group 3 (G3) and group 4 (G4), respectively. The total numbers of miRNAs in each group are 702, 178, 92, 22 for G1, G2, G3 and G4, respectively. Only 2% of total miRNAs have MREs more than 20 (Group G4), whereas ∼70% of the miRNAs were in the group G1 with ≤ 5 MREs. The miRNAs present in G4 are shown in figure 3a, with their number of MREs on 3’UTR written in brackets. The connections in circos plot show the presence of binding sites in each Alu. Non-Alu region is grouped as one and shown at the bottom of the circos. Majority of sites are present in Alus as all the connections from each of the group fall in all the Alu elements. Only ∼ 5% of miRNA binding sites fall in non-Alu regions **(Figure 3a)**.

**Figure 3:**
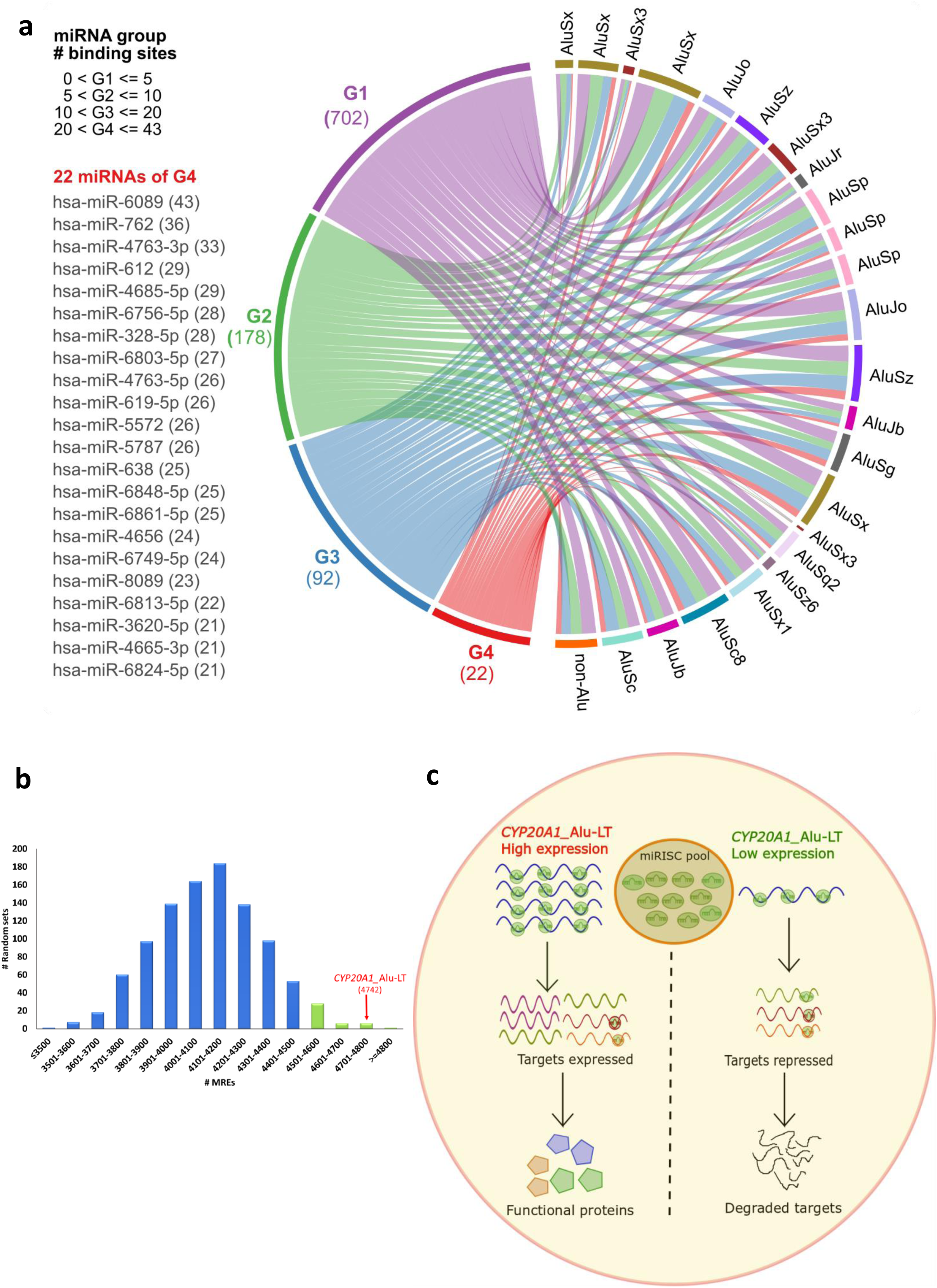
*CYP20A1_Alu-LT* putative lncRNA may act as a miRNA sponge. (a) Circos plot representing the MREs for the 994 miRNAs on *CYP20A1_Alu-LT* 3’UTR. miRNAs are grouped on the basis of number of MREs. 23 Alus in this 3’UTR contribute to 65% of its length and are distributed throughout the UTR. Only 11% of miRNAs have MREs >10 (92 and 22 in G3 and G4, respectively). (b) Distribution of MREs for these 994 miRNAs on 1000 random sets of 23 length and subfamily matched Alu repeats. Only 6 sets contain MREs in range of 4701-4800 suggesting this is a non random phenomenon and MREs are created post Alu exaptations. Highlighted in green are sets with more than 4500 MREs. (c) Proposed model to demonstrate the effect of potential sponge activity of CYP20A1_Alu-LT. In the condition where it is highly expressed, it will recruit multiple miRISC complexes which could relieve the repression of cognate targets leading to their translation; whereas in case of its reduced expression, those miRISC complexes remain free to load on the cognate targets and affect translational repression or promote mRNA degradation. *CYP20A1_Alu-LT* has the potential to sponge multiple miRNAs at the same time thereby regulating a large repertoire of transcripts.

It is plausible that the accumulation of so many Alu-MREs in this 3’UTR has been due to the retrotransposition or recombination of Alus with pre-existing target sites. To test this possibility, we carried out analysis on 1000 sets of 23 Alus taken randomly from the genome with matched length composition and subfamily. We did not observe a similar distribution of Alu-MREs - only 0.5 and 4.2% of these random sets had MREs ≥4742 and 4500, respectively **(Figure 3b)**. This suggests that the chance of Alus having retrotransposed into this 3’UTR with pre-existing MREs is extremely low and these have been created within Alus post exaptation into *CYP20A1_Alu-LT* 3’UTR. One possibility is that accumulation of MREs could potentiate its function as miRNA sponge for a regulatory network **(Figure 3c)**.

### *CYP20A1_Alu-LT* isoform functions as a potential miRNA sponge

To determine if *CYP20A1_Alu-LT* can be a potential miRNA sponge, we characterized this 3’UTR further using bioinformatics and experimental approaches. First, we checked its level using RT-qPCR in both nuclear and cytosolic fractions and found that it is predominantly localized to the cytosol - a feature observed in most sponges **(Figure 4a)**. A sponge RNA also typically contains 4 to 10 low binding energy MREs for a particular miRNA that are separated by a few nucleotides and is generally devoid of destabilizing RNA elements. In *CYP20A1_Alu-LT*, using a stringent cut-off for MRE prediction (binding energy≤ xy425kcal/mol), we observed miRNAs with as many as 43 MREs and binding energy as low as −47kcal/mol. Out of the 994 miRNAs, 140 have ≥10 MREs and are distributed across the length of the UTR **(Figure 3a)**.

**Figure 4:**
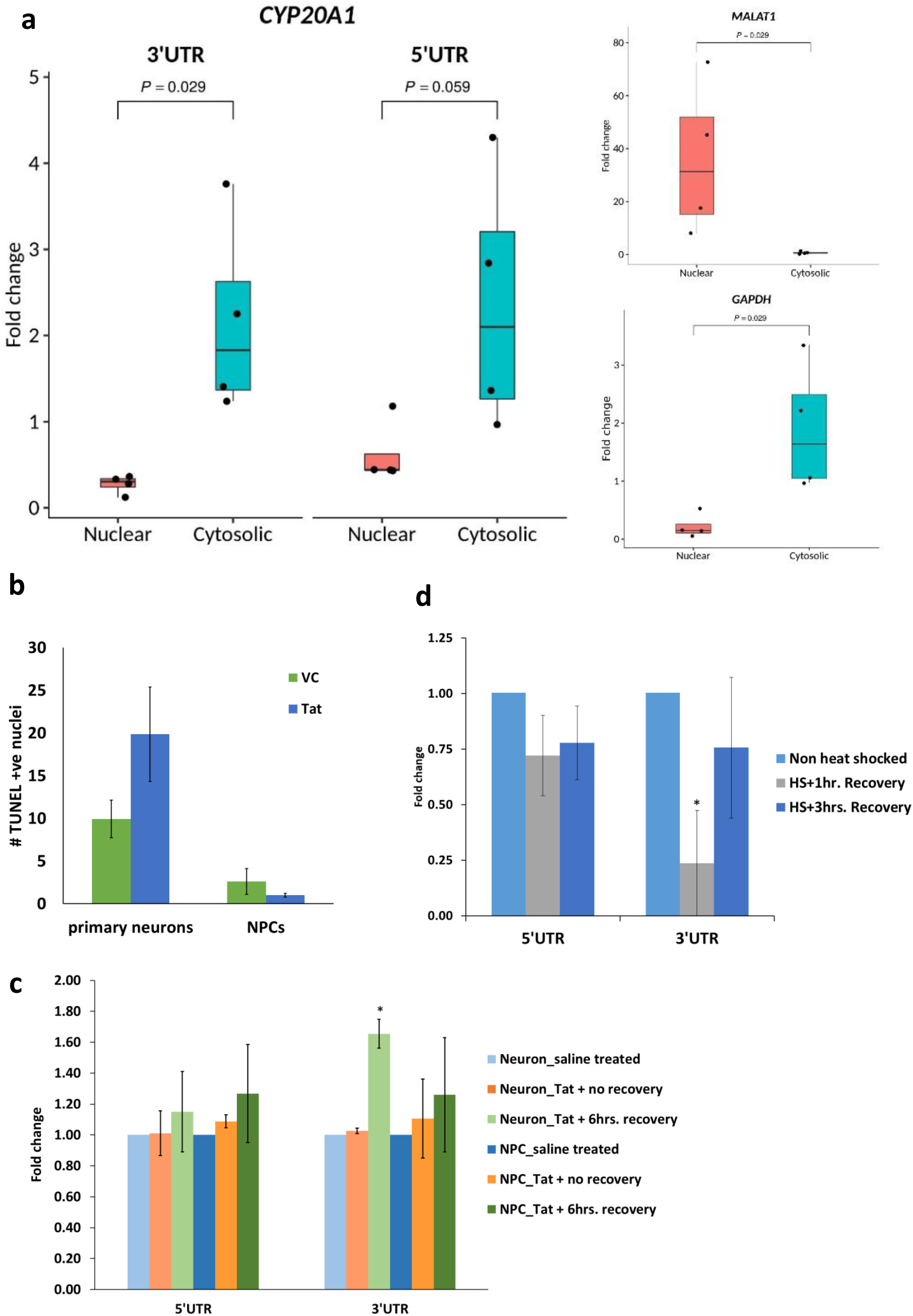
Features of *CYP20A1_Alu-LT* for being a potential sponge RNA. (a) Cytosolic localisation of *CYP20A1_Alu-LT* confirmed by RT-qPCR. Fold change was calculated with respect to total RNA, after internal normalization using the primers against spiked-in control. The error bars represent the SD of four independent experiments and the average of two technical replicates was used for each experiment. Quality controls for assessing the purity of nuclear (*MALAT1*) and cytosolic (*GAPDH*) fractions are shown on the right. The RT-qPCR data were analyzed in accordance with the MIQE guidelines (Bustin et al. 2009) **(Supplementary information, S3)**. (b) Late apoptotic cells in primary neurons and NPCs in response to HIV1-Tat treatment were scored by the number of TUNEL positive nuclei. Tat is neurotoxic and kills ∼50% more neurons compared to the vehicle control (VC i.e., saline), whereas the difference is not statistically significant for NPCs (p-values 0.04 and 0.21 for primary neurons and NPCs, respectively, for Student’s t-test assuming equal variance). The data represents the mean and SD of three independent experiments and >1000 nuclei were scored per condition for each experiment. (c and d) Expression of *CYP20A1_Alu-LT* in response to HIV1-Tat (c) and heat shock (d) treatment was assessed by RT-qPCR using both 5’ and 3’UTR primers. The 3’UTR was found to be upregulated following 6hrs recovery after Tat treatment in neurons (p value = 0.035 *p value <0.05, Student’s t-test), but not in NPCs (p value = 0.348) (c). It was also strongly downregulated in neurons (p value = 0.031) immediately after heat shock (HS+1hr recovery). This difference was not significant during recovery (p value = 0.310; HS+3hrs recovery) (d). In both these cases, the 5’UTR primer exhibits the same trend as the 3’UTR but does not qualify the statistical significance cut-off of p< 0.05. Fold change was calculated with respect to saline (vehicle) treatment, after internal normalization with the geometric mean of *GAPDH, ACTB* and *18S rRNA* in (c) and with respect to non heat shock treated cells, after internal normalization with the geomean of *GAPDH* and *ACTB* in (d). The error bars represent the SD of three independent experiments and the average of 2-3 technical replicates was taken for each experiment.

We next checked for the presence of bulge within the MREs for the 23 prioritized miRNAs **(Table 2)** using miRanda with default parameters. To screen for MREs that would efficiently dock miRNA without degrading the *CYP20A1_Alu-LT* transcript, we used twin criteria – a complete match (2-7) in 6-mer seed site and presence of mismatch or insertion at 9-12 position. 6-mer sites with wobble base pairing were also retained as two wobble-pairs were maximally present in some of the MREs. We found five such sites for miR-6724-5p, two each for miR-1254, miR-4767 and miR-3620-5p and one each for miR-941, miR-4446-3p, miR-296-3p, miR-619-5p, miR-6842-3p and miR-1226-5p **(Table 3)**. At all these sites, we observed insertion in *CYP20A1_Alu-LT*, which suggests the possibility of a bulge formation in the sponge RNA. This can potentially prevent *CYP20A1_Alu-LT* from miRNA directed degradation and increase its efficiency to sequester miRNA molecules.

**Table 2:**
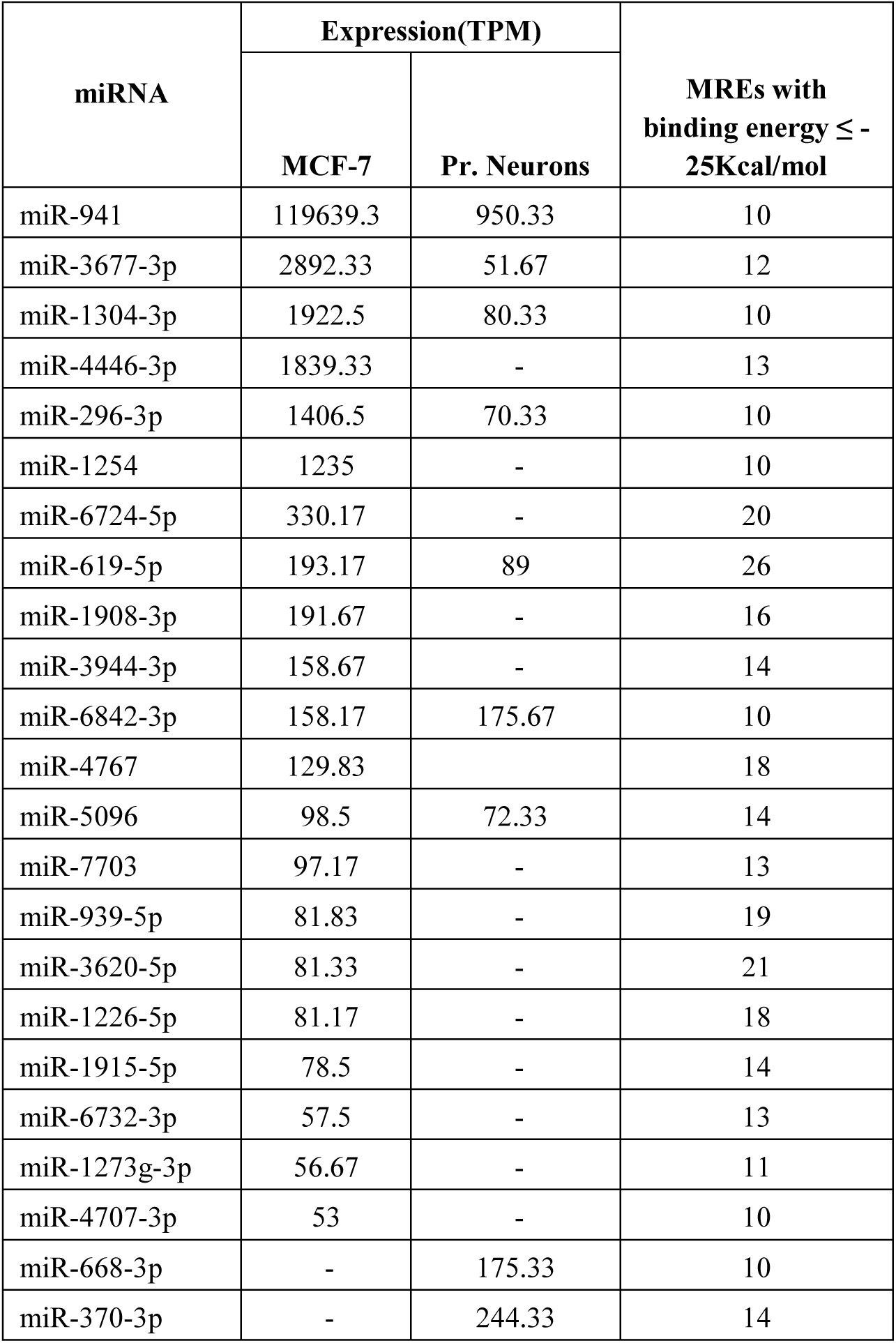
List of 23 prioritized miRNAs. miRNAs were prioritized based on their expression level (≥50 TPM), number of MREs ≥10 with binding energy ≤ 25kcal/mol. * expression values <50 TPM have not been represented.

**Table 3:** Features of MREs with seed site match and presence of bulge.

| miRNA | Total no. of MREs on CYP20A1 3'UTR | Average of Overall complementarity of miRNA with CYP20A1 3'UTR | MRES with seed match(2-7 nt) including wobble | Number of MREs with mismatch or insertion at bulge position | Features of MREs |  |  |
| --- | --- | --- | --- | --- | --- | --- | --- |
|  |  |  |  |  | Forward score | Binding energy | Alignment length |
| miR-941 | 29 | 59.37 | 2 | 1 | 146 | -22.05 | 21 |
| miR-3677-3p | 57 | 54.14 | 3 |  |  |  |  |
| miR-1304-3p | 28 | 59.57 | 12 |  |  |  |  |
| miR-4446-3p | 34 | 60.56 | 3 | 1 | 120 | -27.05 | 22 |
| miR-296-3p | 42 | 54.76 | 5 | 1 | 146 | -25.23 | 25 |
| miR-1254 | 45 | 59.72 | 18 | 2 | 126,128 | -22.05, -23.75 | 25,23 |
| miR-6724-5p | 105 | 44.18 | 19 | 5 | 99,99,103,107, 126 | -21.14, -23.87, -26.94, -22.16, -23.6 | 22,22,22, 21,18 |
| miR-619-5p | 45 | 67.17 | 27 | 1 | 115 | -21.76 | 20 |
| miR-1908-3p | 43 | 52.60 | 1 |  |  |  |  |
| miR-3944-3p | 64 | 54.82 | 23 |  |  |  |  |
| miR-6842-3p | 39 | 59.20 | 11 | 1 | 104 | -24.14 | 29 |
| miR-4767 | 72 | 54.71 | 9 | 2 | 101,124 | -25.3, -23.93 | 27, 16 |
| miR-5096 | 22 | 65.80 | 8 |  |  |  |  |
| miR-7703 | 51 | 57.20 | 4 |  |  |  |  |
| miR-939-5p | 63 | 51.32 | 9 |  |  |  |  |
| miR-3620-5p | 77 | 53.71 | 32 | 2 | 108,130 | -21.24, -22.89 | 22,20 |
| miR-1226-5p | 73 | 56.95 | 6 | 1 | 124 | -26.76 | 29 |
| miR-1915-5p | 40 | 54.88 | 1 |  |  |  |  |
| miR-6732-3p | 35 | 60.37 | 0 |  |  |  |  |
| miR-1273g-3p | 30 | 67.30 | 12 |  |  |  |  |
| miR-4707-3p | 30 | 58.03 | 6 |  |  |  |  |
| miR-668-3p | 38 | 52.86 | 12 |  |  |  |  |
| miR-370-3p | 56 | 56.57 | 2 |  |  |  |  |

### Potential sponge activity of *CYP20A1_Alu-LT* in primary neurons in response to heat shock and HIV1-Tat

To probe if the alteration in *CYP20A1_Alu-LT* level could affect expression of transcripts containing cognate MREs, we looked for conditions where it is likely to be altered. In these conditions the miRNA that target these MREs should also be expressed. We anticipated that in conditions where there is a higher expression of the potential ‘sponge’ (*CYP20A1_Alu-LT*), the abundant MREs would sequester the miRNAs. This should relieve its other cognate targets and we should observe a higher expression of those genes. Whereas in conditions where *CYP20A1_Alu-LT* is downregulated, the miRNA would be free to bind its cognate targets, thereby reducing their expression.

We first queried for the expression of the 994 miRNAs having potential MREs in *CYP20A1_Alu-LT* from miRNA expression profiles available in public datasets. These experiments, mostly microarray based, showed low concordance across replicates and high variability across experiments **(Supplementary information, S1)**. So we tested this experimentally in MCF-7 and primary neurons. Since primary neurons preferentially express longer 3’UTRs, we reasoned it would be an ideal model to study miRNA-mediated regulation events (37,38). We carried out small RNA-seq and using a cut-off of at least 10 MREs on *CYP20A1_Alu-LT* 3’UTR and TPM value of 50, we obtained a set of 21 and 9 miRNAs in MCF-7 and neurons, respectively, of which 7 were common to both **(Table 2)**.

Since *CYP20A1* has been identified as a candidate from a set of Alu exonized genes that map to apoptosis, we asked if this would respond to triggers that induce cell death. HIV1-Tat is a potent neurotoxin that kills ∼50% more neurons compared to the vehicle control **(Figure 4b)**. Upon treating primary human neurons with HIV1 full length Tat protein, followed by 6 hours recovery, *CYP20A1_Alu-LT* was found to be significantly upregulated (1.65 fold). However, progenitor cells, which are immune to Tat **(Figure 4b)**, did not show any such trend **(Figure 4c)**. *CYP20A1_Alu-LT*’s 3’UTR also carries 17 potential binding sites for HSF1, 14 of them within Alus, which show positional conservation in agreement with previous studies (39). This suggests that this transcript may also be amenable to antisense-mediated downregulation during heat shock response as demonstrated by an earlier work from our lab (39). We found *CYP20A1_Alu-LT* to be significantly downregulated (2.68 folds) in primary neurons upon heat shock, followed by 1hr recovery **(Figure 4d)**.

In order to query the expression of the other cognate targets of the 9 prioritized miRNAs in these two conditions, we performed stranded RNA-seq of primary neurons after these treatments. The expression of *CYP20A1_Alu-LT* in RNA-seq showed similar patterns of expression as observed in RT-qPCR, significantly downregulated 2.68 folds (log2FC=-1.42) upon heat shock (HS) recovery and 1.21 folds upregulated (log2FC=0.28) during Tat response. The latter, however, did not cross our stringent statistical significance threshold. Out of the 3876 genes differentially expressed in HS or Tat, 380 exhibit positively correlated expression patterns as *CYP20A1_Alu-LT* **(Figure 5, Supplementary Table S7)**. All of these 380 genes contain at least one MRE for one or more of the 9 prioritized miRNAs and the majority of their MREs are canonical and not Alu-derived. On the contrary, *CYP20A1_Alu-LT* contains a total of 116 MREs for all these 9 miRNAs combined **(Supplementary Table S7)**.

**Figure 5:**
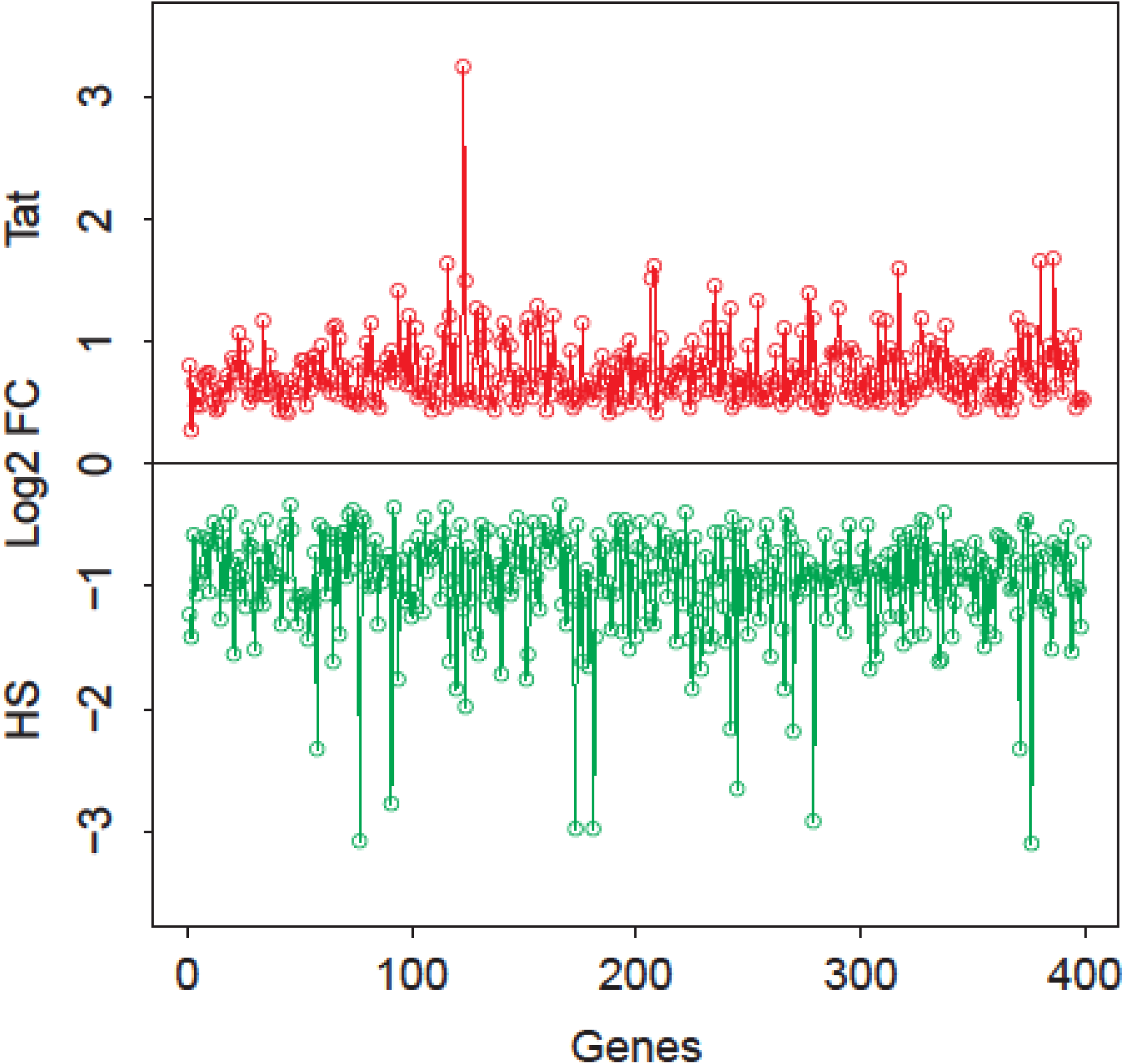
Fold change (Log2FC values) of 380 genes. Figure represents upregulated genes in response to Tat treatment (red) and downregulated during heat shock recovery (green) in primary neurons, resonating with the trend exhibited by *CYP20A1_Alu-LT*. All the transcripts contain one or more MREs for the 9 miRNAs that can be potentially titrated by sponge activity of *CYP20A1_Alu-LT* in neurons. These represent potential cognate targets whose expression can be regulated by *CYP20A1_Alu-LT* perturbation. Genes are plotted in order as **Supplementary table S7**.

Gene Ontology analysis for this gene set using Toppfun revealed enrichment of blood coagulation pathways and neuronal development as major hubs in biological processes. The top five processes were hemostasis (28 genes), axon guidance (25 genes), neutrophil degranulation (23 genes), platelet signaling, activation and aggregation (18 genes) and ECM organization (18 genes). Other processes include mRNA processing and mitochondria translation, metabolism, amino acid and nucleotide synthesis and antigen presentation **(Supplementary Table S8)**.

## Discussion

### Exonized Alus create a unique 3’UTR in *CYP20A1_Alu-LT*

In this study, we highlight a transposable element (TE)-derived putative miRNA sponge that has originated from *CYP20A1* gene with 23 Alus from different subfamilies leading to the formation of a 9kb long 3’UTR. This acts as a substrate for the evolution of thousands of miRNA binding sites. *CYP20A1_Alu-LT* isoform seems to have neo-functionalized from a protein coding locus and its expression is restricted to the higher primates. Our study in primary human neurons suggests that differential expression of this RNA could modulate expression of multiple modules of regulatory network and govern specific outcomes by synchronizing environmental cues.

*CYP20A1* is an orphan member of the cytochrome P450 superfamily of oxidoreductases that metabolize and detoxify drugs (40). It contains unusual heme-binding motifs and is highly expressed in the liver. Efforts to identify its bonafide substrates and inhibitors are underway and some proluciferins have been reported to be its physiological substrates (41). Elevated expression of *CYP20A1* RNA has been reported in splenasthenic syndrome patients as well as blunt trauma patients (42). Besides, high levels of *CYP20A1* mRNA in both unfertilized zebrafish eggs and developing mouse embryos suggest its involvement in early developmental stages (43,44). However, these observations are correlational and it is still contentious whether the gene could function as RNA, a protein or both.

We report both coding as well as non-coding transcripts from the *CYP20A1* locus. While the protein is highly conserved, we observe a novel transcript with a skipped exon that results in an out-of-frame CDS. Further, the RNA isoform has a significantly longer UTR with plausible modulation of post-transcriptional regulatory events. We characterized this isoform using molecular, biochemical, genomics and bioinformatics approaches. It is not only expressed in 75% of single nuclei derived from different layers of the human cerebral cortex but also in rosehip neurons - a highly specialized cell type in humans, suggesting that *CYP20A1_Alu-LT* may play an important, hitherto uncharacterized biological function. This is further corroborated by its presence exclusively in the higher primates, while the protein coding isoform is conserved across the vertebrate phylogeny.

### *CYP20A1_Alu-LT*: a potential miRNA sponge

Among TEs, Alus contain the maximum number of MREs (45). miRNA sponges so far have been mechanistically characterized to contain MREs for a single or a few related miRNA species (46–48). Their involvement in modulating gene expression as well as modifying existing miRNA regulatory networks in a lineage specific manner has been highlighted. For example, miR-1285-1 is processed from an Alu and predominantly targets exonized Alus (45), 3’UTRs of MDM2 and MDM4 harbour Alu-MREs for primate specific miR-661 that provides an additional layer of regulation (49). In patient derived cells from two different cancerous states, it has been shown that elevated levels of free Alu RNA can sequester miR-566 which correlates with disease progression (50). Our earlier work has reported the functional significance of MREs within 3’UTR Alus in fine tuning the p53 regulatory network during stress response (20).

Artificially created sponges with MREs ≥4 have been effectively used in a few studies (51,52) but *CYP20A1_Alu-LT* having ≥10 MREs for as many as 140 miRNAs stands out. Out of the 4742 total MREs more that 80% are within Alu which could potentially participate in miRNA regulatory network. The presence of MREs for miRNAs raises the exciting possibility that it can exert a systemic effect through titrating the cellular levels of many miRNAs simultaneously.

### *CYP20A1_Alu-LT* sponge activity could modulate mRNA-miRNA networks in neuro-coagulopathy

We observe that expression of *CYP20A1_Alu-LT* in human primary neurons is inducible in response to HIV1-Tat and decreased during HS response. Expression of competing endogenous (ce)RNAs, such as sponge RNAs, is also tightly regulated and often, specific to tissue, development stage or stress conditions (53,54). Since sponge RNA could titrate miRISC complexes, their expression could correlate with expression of mRNA having shared targets (51,52) **(Figure 3c)**. We make similar observations in a set of 380 genes which correlate with the expression pattern of *CYP20A1_Alu-LT* i.e., downregulated during heat-shock and upregulated upon Tat treatment. These genes map to processes that are involved in blood coagulation and neuronal pathways. Blood coagulation factors have been reported to affect pathophysiology of CNS via coagulation protein mediated signal transduction (55). Besides coagulation, these proteins interfere with synaptic homeostasis. They also affect neurite outgrowth and morphological changes in neurons, blood brain barrier integrity, ECM stability, ROS generation from astrocytes, secretion of inflammatory cytokines, as well as impair excitability, increase inflammation, mitosis of astrocytes and/or microglia and ultimately, affect neuronal viability. This process is being linked to several neurodegenerative diseases including multiple sclerosis, cancer of the CNS, addiction and mental health (55). Although the exact biological role of *CYP20A1_Alu-LT* remains to be mechanistically elucidated, yet enrichment of coagulation pathways in gene set showing correlated expression with this transcript suggests that it may be involved in fine-tuning inflammatory responses in neuro-coagulopathy, a possibility for future studies.

Exposure to HIV1-Tat is known to cause axonal damage, loss of blood brain barrier integrity, changes in neurite outgrowth, etc. These are mediated by astrocyte activation, inflammatory cytokine expression, inducing mitochondrial injury and rearrangement of microtubules. The set of 380 genes which correlate with the expression pattern of *CYP20A1_Alu-LT* were also enriched in similar pathways like axon guidance, hemostasis, platelet activation and aggregation, ECM organization, regulation of actin cytoskeleton, antigen presentation, Golgi to ER transport and mitochondrial translation. In the light of our observations, it is possible that the changes observed upon Tat exposure could partly be mediated and synergised by the sponging effect of *CYP20A1_Alu-LT*. Upon activation by Tat, the sponge could titrate out the miRNA that target the 380 genes and hence modulate all the pathways simultaneously. Validation of *CYP20A1_Alu-LT* in the context of neuronal damage might shed further light on this plausible involvement. It could also play a role in normal neuronal functions through fine tuning expression in NPCs during neurogenesis, neuronal migration during differentiation, etc.

### Future directions and alternate possibilities

Multiple exonized Alus in *CYP20A1_Alu-LT* 3’UTR can facilitate secondary structure and lead to altered bioavailability for MREs. Future studies on secondary structure simulations would allow us to assess the availability and accessibility of these MREs. Closely spaced sites of same or different miRNAs can cooperatively sequester multiple miRNAs thus making the process fast and leading to robust outcomes (56). With multiple miRNAs that target members of a co-regulated network, differential expression of a sponge could exhibit a systemic response. Also, this could work as an effective regulatory switch for a faster response or return to baseline. We have not yet looked at its turnover rates; however multiple possibilities exist such that it could be transiently induced in response to specific environmental cues, regulated through a negative feedback, cleared via transcriptional shutdown by certain miRNAs or has an inherently shorter half-life due to a rapid turnover. Other possibilities such as independent transcription events from this UTR or additional polyadenylation sites also cannot be ruled out. During the course of this study, we noticed reads mapping beyond the longest 3’UTR annotated in UCSC **(Supplementary Figure 5)** raising the possibility that isoforms with even longer 3’UTRs may be transcribed from this locus. Beyond its role as miRNA sponge, this UTR can also be involved in sequestering RNA binding proteins and deplete their cellular reserves, thereby indirectly affecting other genes (57). Further, all the 23 Alus in this 3’UTR have been shown to harbour antisense transcripts and are substrate for A-to-I RNA editing (19). These events, though dynamic, could further disrupt or create new miRNA binding sites from existing MREs, thereby increasing the regulatory repertoire. Events such as this have been reported in miR-513 and miR-769 that target 3’UTR of DNA fragmentation factor alpha gene in an adenosine deamination dependent manner (58). Since A-to-I editing events are preponderant in the brain, these editing events could further contribute to phylogenetic novelties (59,60).

## Conclusion

This study adds to the growing repertoire of regulatory functions of Alu in the human transcriptome. In this study, we provide a novel dimension of its regulatory potential -that of creation of a miRNA sponge through Alu exaptations in the 3’UTR regions. *CYP20A1* provides an interesting model for studying Alu derived novel transcripts that can function as ceRNAs and co-regulate multiple genes in a network or cellular process. Thus, the addition of a lineage specific sponge could be a top-up on existing networks that modulate intermediate phenotypes

such as neuro-coagulation. These could act as regulatory switches and in response to biological cues rapidly release or sequester miRNAs to govern specific cellular outcome.

## Materials and Methods

### Bioinformatics

#### Characterization of a novel transcript isoform of CYP20A1

Extensive annotation of different transcript isoforms of CYP20A1 was carried out using Ensembl, NCBI and UCSC. Details are provided in **Supplementary information S1**.

#### Length comparison of the 3’UTR of CYP20A1_Alu-LT with other 3’UTRs at genome-wide scale

The coordinates for human transcripts (NM and XM IDs) were downloaded from NCBI RefSeq version 74 (hg38). For every gene, only the longest 3’UTR was considered. The summary statistics for size distribution were calculated using R scripts.

#### DNA conservation analysis

DNA sequence conservation across different species was checked with UCSC genome browser using multiple alignment across 20 species generated by multiz (61). Both gaps as well as unaligned sequences were treated as ‘missing’ data.

#### Protein conservation analysis

CYP20A1 protein sequences from different species were taken from the top hits obtained in NCBI pBLAST by using the human protein as a reference. Multiple sequence alignment was performed using Clustal Omega (O 1.2.2). As described in Gautam et al. 2015 Ka/Ks ratio was calculated (see Supplementary information, S1 for details) (62).

#### *CYP20A1_Alu-LT* expression in non-human primates

We used publicly available chimp and macaque RNA-seq datasets from GEO [GSM1432846, 55, 65 (SRR1510158, 167, 177); GSM2265102, 4, 6 (SRR4012405, 08, 09, 13)]. Reads were mapped to both human and chimp/macaque 3’UTR to increase fidelity and mapping on housekeeping genes like *ACTB, GAPDH* and *EIF4A2* was also checked to control for data quality and mapping parameters. To query more expression datasets, we took advantage of the sequence differences in this transcript due to skipping of sixth exon. We performed BLAST against human datasets in SRA using a 289bp sequence reconstructed by joining exon 5 and 7. The hits were reconfirmed by alignment of reads to the 3’UTR.

RNA-Seq Reads from non-human primate reference transcriptome mapped on hg19 were exported as UCSC genome browser tracks. We additionally incorporated the stranded RNA-seq data generated as a part of this study to compare the expression level of this transcript between human and other non-human primates.

#### miRNA target prediction in CYP20A1_Alu-LT

miRNA target sites (MREs) on *CYP20A1* 3’UTR were predicted using miRanda (version 3.3a) (36), with the parameters set as follows: score threshold(-sc): 100, gap opening penalty(-go): −8, gap extension penalty(-ge): −2, binding energy(-en): −25kcal/mol., ‘strict’ (i.e., G:U pairs and gaps were not tolerated in the seed region). miRanda uses miRBase (which contains ∼2500 miRNAs) for annotation. For bulge analysis, target prediction for 23 miRNAs on *CYP20A1* 3’UTR was performed using miRanda offline version 3.2a with default parameters (gap opening penalty= −8, gap extension= −2, score threshold= 50, energy threshold= −20kcal/mol, scaling parameter= 4).

#### RNA-seq

Fastq files were checked using FASTQC and overall Q score was >20 with no adapter contamination. Overrepresented sequences were not removed. Reads were mapped on hg38 using Tophat, followed by isoform quantification (Cufflinks) and collation (Cuffmerge). Overall read mapping rate was between 59-86.3% and concordant pair alignment ranged between 53.1 and 81.1%. Cuffdiff was used to calculate the differential expression (D.E.; calculated for each experimental condition against untreated). Summary of sample-wise RNAseq data is provided in **supplementary information (S3)**.

#### Small RNA-seq

The data were quality checked using FastQC (version 0.11.2), followed by adapter trimming by cutadapt (version 1.18) and reads were not discarded. As expected, around 95% of the adapter trimming events happened at the 3’end of the reads. Filtering based on length and quality were carried out by cutadapt; Q30 reads with sequence length >15 but < 35nt were retained for mapping. Nearly 80% of the reads were retained after these filtering steps. Size distribution of the reads and k-mer position were (21-25, 28-32) and 26-28 respectively. Subsequently, these reads were mapped onto hg38 using Bowtie2. On average, 61% of the reads were uniquely mapped. miRDeep2 was run to obtain the read counts as TPM. Summary of sample-wise small RNAseq data is provided in supplementary information (S3).

### Experimental

#### Expression analysis of *CYP20A1_Alu-LT* across diverse cell lines

##### RNA isolation, cDNA synthesis and RT-qPCR

Total RNA was isolated using TRIzol (Ambion, Cat. No. 15596-026) as per manufacturer’s protocol and its integrity was checked on 1% agarose gel followed by Nanodrop quantification (ND1000, Nanodrop technologies, USA). cDNA was prepared from oligo(dT)-primed DNase-treated RNA (Invitrogen, Cat. No. AM1907) and SuperScript III RT (Invitrogen, Cat. No. 18080-044). RNA template was digested from the cDNA using 2 units of E. coli RNaseH (Invitrogen, Cat. No. 18021071). Primers were designed using Primer3 (version 4.0.0) and were synthesized by Sigma **(Supplementary information, S2)**. To ensure there was no spurious amplification, we designed two pairs of overlapping primers both on the 5’ as well as 3’ends of our transcript of interest and included ‘minus-RT’ controls in every reaction. Additionally, we sequenced three amplicons (1, 5 and 10) to check the specificity of amplification **(Supplementary information, S1)**. BLASTN (NCBI; 2.4.0+) against the corresponding *in silico* predicted amplicons had revealed >95% sequence identity with an average query cover of 90%; BLAT against the whole genome (hg38) gave CYP20A1 as the top hit in every case. RT-qPCR was performed using 2X SYBR Green I master mix (Kapa Biosystems, Cat. No. KK3605) and the reaction was carried out in Roche LightCycler 480 (USA) **(Supplementary information, S1)** Melting curves were confirmed to contain a single peak and the fold change was calculated by ^ΔΔ^Ct method. MIQE guidelines were followed for data analysis.

##### 3’RACE for mapping the full length transcripts

cDNA for 3’RACE was prepared using RLM-RACE kit (Ambion, Cat. No. AM1700) with 1µg MCF-7 total RNA as per the manufacturer’s recommendation. Nested PCR was performed with FP10 and an internal primer using the amplicon produced by FP9 and external primer. The product of this nested PCR was electrophoresed on 2% agarose gel and four major bands were observed, which were gel eluted using Qiaquick gel extraction kit (Qiagen, Cat. No. 28704) and subsequently sequenced. Details of the results are provided in the **Supplementary information, S3**.

##### Cell culture studies

MCF-7 cell line was procured from National Centre for Cell Sciences, (Pune, India) and cultured in GlutaMax-DMEM high glucose (4.5gm/l) (Gibco, Cat. No. 10569044) supplemented with 10% heat inactivated FBS (Gibco, Cat. No. 10082147), HEPES (Gibco, Cat. No. 11560496) and 1X antibiotic-antimycotic (Gibco, Cat. No. 15240096). The culture was maintained at 70-80% confluency at 37°C, 5% CO^2^. Cell line lineage was confirmed by STR profiling and cells were routinely screened for any contamination **(Supplementary information, S1)**.

Primary human neuron and astrocyte cultures comply with the guidelines approved by the Institutional Human Ethics Committee of NBRC as well as the Stem Cell and Research Committee of the Indian Council of Medical Research (ICMR) (Fatima et al. 2017). Briefly, neural progenitor cells (NPCs) derived from the telencephalon region of a 10-15 week old aborted foetus were isolated, suspended into single cells and plated on poly-D-lysine (Sigma, Cat. No. P7886) coated flasks. The cells were maintained in neurobasal media (Gibco, Cat. No. 21103049) containing N2 supplement (Gibco, Cat. No. 17502048), Neural Survival Factor 1 (Lonza, Cat. No. CC-4323), 25ng/ml bovine fibroblast growth factor (bFGF) (Sigma, Cat. No. F0291), 20ng/ml human epidermal growth factor (hEGF) (Sigma, Cat. No.E9644) and allowed to proliferate over one or two passages. The stemness of NPCs was functionally assayed by - i) formation of neurospheres, and ii) ability to differentiate into neurons or astrocytes. Additionally, NPCs were also checked for the presence of specific markers like Nestin. For commitment to the neuronal lineage, NPCs were starved of bFGF and EGF; with 10ng/ml each of PDGF (Sigma, Cat. No. P3326) and BDNF (Sigma, Cat. No. B3795) added to the media cocktail. Differentiation of NPCs to astrocytes required Minimum Essential Medium (MEM) (Sigma, Cat. No. M0268-10x) supplemented with 10% FBS. The process of neuronal differentiation completes in exactly 21 days; our experiments were completed within a week post-differentiation. Differentiated cultures of primary neurons and astrocytes were also checked for specific markers by immunostaining to determine the efficiency of the differentiation process **(Supplementary Figure S6)**.

##### Nuclear -cytosolic localization of *CYP20A1_Alu-LT*

Nuclear and cytosolic RNA were isolated using PARIS kit (Ambion, Cat. No. AM1921) as per manufacturer’s protocol. Briefly, nearly 10 million cells were resuspended in fractionation buffer, incubated on ice and centrifuged at 4°C to separate the nuclear and cytosolic fractions. The nuclear pellet was additionally treated with cell disruption buffer before mixing with the 2X lysis/binding solution and absolute ethanol and passing through a column. The RNA was subsequently eluted in hot elution buffer; quantified using Nanodrop and its integrity was checked on 1% agarose gel. Nuclear RNA contains an additional hnRNA band above the 28S rRNA band and is usually of lower yield than cytosolic RNA. RT-qPCR was done as described earlier using gene specific primers from 5’UTR and 3’UTR region.

##### Induction of Stress

Cells were gently washed once with 1X PBS and fresh media was replenished before treatment for accurate quantification of stress response. Heat shock was given at 45°C (±0.2) for 30 min in a water bath. Subsequent to the treatment, cells were transferred to 37°C/5% CO^2^ for recovery and harvested after 1hr, 3hrs and 24hrs. For Tat treatment, full-length lyophilized recombinant HIV1 Tat protein was purchased from ImmunoDX, LLC (Woburn, Massachusetts, USA) and reconstituted in saline. The dosage for treatment was determined by drawing a ‘kill curve’ using graded dose of Tat on neurons **(Supplementary Figure S7)**. Treatment was performed for 6hrs with 100ng/ml Tat and cells were either harvested just after the treatment or allowed to recover at 37°C and 5% CO^2^ for another 6hrs prior to harvesting.

##### TUNEL Assay

The assay was performed with in situ Cell Death Detection kit, TMR red (Millipore Sigma, Cat. No. 12156792910). Nearly 20,000 cells were seeded per well (on coverslips) in 12-well plates. Post Tat treatment, cells were washed once with 1X PBS and fixed with 4% PFA, followed by three washes with 1X PBS, permeabilization and blocking with 4% BSA containing 0.5% Triton-X 100, incubation with TdT for 1hr in the dark and three washes with 1X PBS. Coverslips were then mounted on clean glass slides using hardset mounting media containing DAPI (Vectashield, Cat. No. H-1500). Six to eight random fields were imaged for each experimental group using AxioImager, Z1 microscope (Carl Zeiss, Germany). Fixed cells treated with 2 units of DNaseI (for 10 minutes at RT, followed by the addition of EDTA to stop the reaction) were used as a positive control in this experiment. TUNEL positive nuclei were scored using ImageJ software (NIH, USA). Minimum of 1000 cells were scored for each replicate.

##### RNA sequencing and small RNA sequencing

Detailed methods for library preparation for RNA-seq and small RNA-seq are provided in supplementary information (S1). Briefly, libraries for RNA-seq were made using 500ng of total RNA per sample and three biological replicates were taken per experimental condition. Libraries were prepared following Illumina’s TruSeq stranded total RNA protocol. The final libraries were pooled, diluted and denatured to a final concentration of 8pM. Clusters were generated using TruSeq PE cluster kit V3-cBOT-HS on cBot system, followed by paired end sequencing on HiSeq2000 using TruSeq SBS kit V3-HS (200 cycles). Libraries for small RNA sequencing were prepared using Illumina’s TruSeq small RNA library preparation kit from 1µg total RNA. The libraries were normalized to 2nM, denatured and subjected to cluster generation on cBot using TruSeq SR cluster kit v3-cBOT-HS. Single read sequencing was performed on HiSeq2000 using TruSeq SBS kit v3-HS (50 cycles).

## Declarations

### Ethics approval and consent to participate

Not applicable

### Consent for publication

Not applicable

## Acknowledgements

The authors acknowledge Chitra Mohindar Singh Singal, NBRC for her help with the TUNEL assay, Parashar Dhapola for his help with data visualization in the initial phase of this work, Dr. Amit Chaurasia for Jukes Cantor divergence analysis and Drs. Debojyoti Chakraborty and Rakesh Dey for providing reagents and many fruitful discussions. The authors would also like to acknowledge Mr.Raghunandanan MV and Mr. Amit Khulve at IT division CSIR-IGIB for their constant help and support for data upload in GEO.

## Availability of data and materials

1. The raw data for small RNA and mRNA sequencing generated in this study have been submitted to GEO (GSE132447).
2. dbGap link for the human temporal cortex (MTG) raw sequence reads https://www.ncbi.nlm.nih.gov/projects/gap/cgi-bin/study.cgi?study_id=phs001790.v1.p1

## Authors contribution

MM and AB designed the study and co-wrote the manuscript along with KS and RP. AB performed conservation analyses, miRNA target prediction, cell culture and molecular biology experiments and helped in RNA seq data analysis. VJ analyzed mRNA and small RNA seq data, ran miRNA target prediction on miRanda and helped in data visualization. KS performed molecular biology experiments, ran miRNA bulge analysis with DD and contributed to improving data visualization along with RK. MF carried out primary human neuron and NPC culture under the supervision of PS. DS performed some of the cellular assays. GC prepared NGS libraries and carried out sequencing, assisted by AB and KS. TEB analyzed the small RNA-seq data. BP contributed reagents, helped in troubleshooting experiments and provided critical inputs. MM supervised the overall study.

