## Supplementary Information for "Origin of a novel *CYP20A1* transcript isoform through multiple Alu exaptations creates a potential miRNA sponge"

#### S1- Materials and methods

##### Mapping 319 genes onto the apoptosis pathway

A transcriptome-wide scan for the co-occurrence of Alu exonization, editing and antisense at the single-exon level had revealed a set of 319 genes, enriched for apoptosis (Mandal et al. 2013). Because enrichment analyses are heavily confounded by background data used for normalization, we found it prudent to ask how many of these 319 genes are actually involved in apoptotic cell death and where in the apoptosis pathway they mapped. To begin with, we listed all the important steps in this pathway: arrest of division is usually the first cellular response to any apoptotic trigger, which is followed by cessation of DNA damage repair (DDR), initiation of p53 signaling and polymerase halt. At the later stages of apoptosis, endonucleases are activated and nuclear DNA fragmentation occurs, mitochondrial ROS may be generated and many proteins are ubiquitinylated and marked for proteasomal degradation. The final stages are marked by blebbing and membrane flipping to attract phagocytes to engulf the dying cell. A flow diagram was constructed incorporating all the nodes and pathways involved in apoptosis. We performed an extensive, systematic manual curation of these 319 genes using published literature and other web resources with the intent of binning them in the major events of apoptosis listed above. We asked i.) which of these genes are involved in apoptosis, and ii.) for those that are involved, where in the pathway they occur. We mapped all those genes which interact with known apoptotic genes (like *BCL2* family members) or contain known motifs (e.g., WD repeats), or are a part of apoptotic signaling pathway, or have apoptosis (or any of its sub-readouts) as the major phenotype upon their perturbation. Many of the genes that we mapped are involved in cell division, DNA replication and ubiquitinylation; perturbations in these genes often lead to cell cycle arrest, impaired DDR and protein misfolding/ degradation, respectively. Many genes are involved in multiple, interlinked functions but we had mapped them uniquely. Mapping was based on the major function of the gene, its mutant phenotype or the function affected by perturbation (knockdown, knock-out or over-expression), disease association and evolutionary conservation (wherever orthologues could be reliably ascertained). To check the accuracy of our curation and mapping, we used GeneMANIA (attributes: physical and genetic interactions, co-localization, same biological pathway) to fetch the top 100 interacting partners of our 91 apoptosis genes. There was no overlap between these interacting partners and the 228 genes that we have not mapped to apoptosis, reassuring us that our mapping stringency has been optimal.

##### Disparities in *CYP20A1* annotation across different portals

There exists a huge disparity in the annotation of *CYP20A1* transcript isoforms in different databases. There are 10, 9 and 4 annotated isoforms in Ensembl, NCBI and UCSC, respectively. All the four UCSC isoforms (GENCODE v24) are mapped onto a single RefSeq transcript NM_177538. Among the 10 isoforms listed in Ensembl, three are targeted for NMD and another three are processed transcripts. In NCBI, out of the 9 transcripts, two are miscellaneous RNAs, six are predicted protein-coding and only one is experimentally validated (NM_177538.2), which is the largest isoform and also annotated as the principal isoform (by APPRIS). It matches with the longest isoform on UCSC in terms of length (10.94kb) as well. Ensembl previously mapped it to ENST00000611416.4 (which has been removed from the current build - release 94) but currently maps this onto its transcript which is just 1949bp long. Notably, the size of the CYP20A1 protein remains the same (462aa) on every portal.

These annotations are based on microarray or RNA-seq data, both of which have their own limitations. Array imputes the expression based on one or few probes and cross hybridization is a potential confound. RNA-seq gives an FPKM value for a transcript regardless of the reads aligning over the entire length of the transcript. The scenario is further complicated because 65% of the long-3'UTR of *CYP20A1*(*CYP20A1*_Alu-LT) is made up of Alu repeats which is typically not captured by array probes and where sequencing reads do not map uniquely. Also, from these data, one cannot discriminate if the full length 3'UTR containing isoform is expressed as a single transcript or as multiple short RNAs.

##### Confirmation of *CYP20A1_Alu-LT* by Sanger sequencing

The sequencing reaction was carried out using Big dye Terminator v3.1 cycle sequencing kit (ABI, 4337454) in 10µl volume (containing 2.5µl purified DNA, 0.8µl sequencing reaction mix, 2µl 5X dilution buffer and 0.6µl forward/ reverse primer) with the following cycling conditions - 3 mins at 95°C, 40 cycles of (10 sec at 95°C, 10 sec at 55°C, 4 mins at 60°C) and 10 mins at 4°C. Subsequently, the PCR product was purified by mixing with 12µl of 125mM EDTA (pH 8.0) and incubating at RT for 5 mins. 50µl of absolute ethanol and 2µl of 3M NaOAc (pH 4.8) were then added, incubated at RT for 10 mins and centrifuged at 3800rpm for 30 mins, followed by invert spin at <300rpm to discard the supernatant. The pellet was washed twice with 100µl of 70% ethanol at 4000rpm for 15 mins and supernatant was discarded by invert spin. The pellet was air dried, dissolved in 12µl of Hi-Di formamide (Thermo fisher, 4311320), denatured at 95°C for 5 mins followed by snapchill, and linked to ABI 3130xl sequencer. Base calling was carried out using sequencing analysis software (v5.3.1) (ABI, US) and sequence was analyzed using Chromas v2.6.5 (Technelysium, Australia).

##### miRNA expression in MCF-7 cells

We wanted to use publicly available expression data to find out which of the 994 miRNAs with predicted MREs on *CYP20A1*_Alu-LT 3'UTR, are expressed in our study system. In order to reduce false positives, we decided to take a consensus of miRNAs found to be expressed in MCF-7 in both microarray (doi:10.1371/journal.pone.0049067) as well as NGS (GSE68246) datasets. In the microarray data, 104 out of our 994 miRNAs of interest were expressed (averaged signal range 7.6 to 974.63). miRNome data of MCF-7 monolayer culture (using Illumina TruSeq small RNA on HiSeq2000) revealed the expression of 102 out of our 994 miRNAs.
Although we found around 100 (out of 994) miRNAs to be expressed in each of these data yet there was absolutely no overlap between the two sets.

##### Ka/Ks calculation

Protein sequence IDs of CYP20A1 ortholog for each species was identified as the first hit of NCBI pBLAST using human NP-ID as the query term. The NM-IDs of corresponding mRNAs were also obtained from NCBI and sequences were fetched for these NP and NM-IDs. For a pair of orthologous genes, the protein-coding DNA alignment between human and each of the other species was constructed using ParaAT (Zhang et al. 2012). The protein sequence information was used to match each triplet codon with its respective amino acid. Aligned coding DNA between a pair (human vs. another species) was provided as an input to KaKs_Calculator (Zhang et al. 2006). Model averaging (MA), which takes an average of substitution rates of seven models, was used as a maximum likelihood method to calculate the Ka/Ks value.

##### RT-qPCR

RT-qPCR was performed in 10µl volume using 5µl 2X SYBR Green I master mix (Kappa; KK3605), 0.2µl each of 10µM forward and reverse primers and cDNA (stock: ~80ng/µl RNA equivalent, diluted 1:40 in the final reaction). The reaction was carried out in Roche Light Cycler 480 (USA) and the cycling conditions were: 95°C for 3', 45 cycles of (95°C for 30", 58°C for 30", 72°C for 40"), 72°C for 3'; followed by melting: 95°C for 5", 65°C for 1'. Melting curves were confirmed to contain a single peak and the fold change was calculated by ΔΔCt method. MIQE guidelines were followed for data analysis.

**Authentication of MCF- 7 Cell lines**

We had authenticated the cell line identity by STR profiling. The cells were routinely screened for mycoplasma using MycoAlert Mycoplasma detection kit (Lonza, LT07-218.) as per manufacturer's protocol and confirmed to be free from contamination. Additionally, 16S rRNA PCR was performed using both genomic DNA from cells and cell-free DNA from spent media to confirm the absence of bacterial contamination.

**Library preparation, RNA sequencing**

Libraries were prepared following Illumina's TruSeq stranded total RNA kit (Illumina, RS-122-2202) protocol as per manufacturer's recommendations, using three biological replicates per experimental condition. Integrity of the RNA samples was checked on Agilent 2100 Bioanalyzer (Agilent RNA 6000 Nano Kit); RIN values ranged from 8.6-9.6 for our samples. 500ng of total RNA was taken in a volume of 10µl and ribosomal RNA was removed using Ribozero depletion kit. Briefly, total RNA was denatured at 68°C for 5', followed by removal of rRNA using magnetic beads, subsequent clean up with RNAClean XP beads (Beckman Coulter, A63987), two consecutive washes in freshly prepared 70% ethanol and then elution (by heating at 94°C for 8'; to fragment and prime the RNA for cDNA synthesis). First strand synthesis was carried out with random hexamers using the following program (25°C for 10', 42°C and 70°C for 15' in each). This was followed by second strand cDNA synthesis at 16°C for 1hr for elimination of the template RNA, replacement of dUTP by dTTP and to finally produce blunt-ended cDNA, which was then cleaned up using AMPure XP beads (Beckman Coulter, A63881) and two washes in 80% ethanol. Next, the 3'ends of these blunt cDNA molecules were adenylated to create a single 'A' overhang (incubation at 37°C for 30', followed by 70°C for 5') such that these can form complementary base pairs with the 'T's on the adapter ends and also do not concatenate with each other to form chimera during adapter ligation. Then, adapters were ligated (30°C for 10') at both the ends of these double-stranded cDNA molecules so that these can hybridize to the probes on the sequencing flow-cell. The fragments with adapter ligated on both ends (libraries) were enriched and amplified using a 15-cycle PCR (the PCR primer cocktail anneal to both ends of the adapters; PCR conditions: 98°C for 30", 15 cycles of (98°C for 10", 60°C for 30", 72°C for 30"), final extension 72°C for 5'). This was followed by their clean up using AMPure XP beads and two washes in 80% ethanol. The libraries were quantified using Qubit fluorometer (Invitrogen) and subsequently validated on Bioanalyzer (Agilent DNA 1000 kit) to confirm the insert size. The size of the libraries was in the range of 260-280bp. This was followed by normalization, dilution of the libraries to 10nM and pooling 3 samples together. The 10nM libraries were denatured using NaOH, diluted to 8pM and clusters were generated on a paired end flowcell (1 pool/lane) using TruSeq PE cluster kit v3-cBOT-HS (Illumina, PE-401-3001) on cBot system. Paired end sequencing was carried out on HiSeq 2000 using TruSeq SBS kit v3-HS (200 cycles) (Illumina, FC-401-3001).

**Library preparation Small RNA sequencing s**

cDNA libraries were prepared as per manufacturer's protocol using TruSeq small RNA library prep kit (Illumina, RS-200-0012, RS-200-0024) from 1µg total RNA (RIN ≥8.5). Briefly, 3' and 5' adapters were ligated, followed by enrichment of RNA fragments with adapter molecules on both ends. Reverse transcription followed by amplification using primers that anneal to the adapter ends created the cDNA libraries which was then purified and size selected (~147nt) to enrich for small RNAs by running on 6% native PAGE. Subsequently, the libraries were concentrated by ethanol precipitation and quality checked on Agilent technologies 2100 Bioanalyzer (USA) using DNA high sensitivity chips (Agilent, 5067-4626). Using 10mM Tris-HCl (pH 8.5) libraries were normalized to 2nM, denatured and applied to cBot for cluster generation (Illumina, GD-401-3001), which were then sequenced on HiSeq2000 (Illumina, California, USA) using TruSeq SBS kit (Illumina, FC-401-3002) for 50 cycles.

**Immunostaining**

Cells were seeded on coverslips and cultured in 4-well plates for this experiment. Spent media was discarded and cells were briefly washed with 1X PBS (to remove residual media completely). The cells were then fixed with 4% PFA (freshly prepared in 1X PBS) for 20 minutes, followed by 3 washes in 1X PBS (5 minutes each) to completely remove traces of PFA. Blocking was done with 4% BSA containing 0.5% Triton-X (to permeabilize the cells) for 1hr with gentle rocking. Incubation in primary antibody (MAP2, Tuj1, NeuN, Nestin, GFAP; all diluted 1:200 in 0.1% BSA) was done overnight at 4°C on a rocker, except GFAP (1hr at RT), followed by 3 washes in 1X PBS (5 minutes each). Incubation with fluorescently labeled secondary antibody (diluted 1:1000 in 0.1% BSA) was performed for 1hr in the dark with shaking. This was followed by five washes in 1X PBS (5 minutes per wash) on a rocker. The coverslips were mounted in DAPI hardset (Vectashield, USA) on clean glass slides. Images were captured from 7-10 random fields using AxioImager.Z1 microscope (Carl Zeiss, Germany).

**MTT assay**

~10,000 cells were seeded per well in 96 well plate. Post treatment, freshly prepared MTT (HiMedia, TC191) solution (in 1X PBS) was added to the cells at a final concentration of 0.5mg/ml. Cells were incubated for 3hrs at 37°C in dark and then checked for formazan crystals under the microscope. After removing the solution, the formazan crystals were dissolved in 100µl DMSO (Sigma, D2650) at RT for 5-10 minutes and then absorbance was acquired at 560nm on Tecan.

#### Supplementary Information S2

#### Primers used in this study

| Oligo Name | Sequence (5'→3') | Length (nt) | Tm (°C) | GC% | Amplicon size (bp) | |
|---|---|---|---|---|---|---|
| CYP20A1_5'UTR_FP1 | TGTCTGAAATCGTGTGCAC | 19 | 56.8 | 47.4 | 222 | used as the "5'UTR primer" |
| CYP20A1_5'UTR_RP1 | CCAATTTAAGGCATAGCGTG | 20 | 55.7 | 45.0 | | |
| CYP20A1_5'UTR_FP2 | ACAGAGCACTACTAACTCCT | 20 | 55.5 | 45.0 | 214 | |
| CYP20A1_5'UTR_RP2 | CAGAGGAAAGAGCAATGGAT | 20 | 55.7 | 45.0 | | |
| CYP20A1_CDS_FP3a | TCACTGTTGATGAGAATGGG | 20 | 55.6 | 45.0 | 655 | |
| CYP20A1_CDS_FP3b | GTATTGGTGAAGAGACTGCA | 20 | 55.7 | 45.0 | 470 | |
| CYP20A1_3'UTR_RP3 | ACAAACATGATTCCCCAACT | 20 | 55.7 | 40.0 | | |
| CYP20A1_3'UTR_FP4 | TCCCTTCCCACTTATTTTCT | 20 | 54.1 | 40.0 | 166 | |
| CYP20A1_3'UTR_RP4 | TTGCTTTAAACTCCACCTCT | 20 | 55.1 | 40.0 | | |
| CYP20A1_3'UTR_FP5 | ACCAAGTGTCAGATCAGATG | 20 | 55.4 | 45.0 | 177 | |
| CYP20A1_3'UTR_RP5 | GCTCTCTTTTCCTTGATTCC | 20 | 54.6 | 45.0 | | |
| CYP20A1_3'UTR_FP6 | GAGAGCATAGAGGAATCAGC | 20 | 55.7 | 50.0 | 216 | |
| CYP20A1_3'UTR_RP6 | GTGAAGTGGCCAAAATTTGA | 20 | 55.5 | 40.0 | | |
| CYP20A1_3'UTR_FP7 | TTGTCACTCTGAATGTAGGC | 20 | 55.7 | 45.0 | 207 | |
| CYP20A1_3'UTR_RP7 | AAAATGAGCTGGGTGTGATG | 20 | 56.9 | 45.0 | | |
| CYP20A1_3'UTR_FP8 | GCCACTACACTCAGCTAATT | 20 | 55.7 | 45.0 | 238 | |
| CYP20A1_3'UTR_RP8 | GTTGATTGAGCCCTCCTTAA | 20 | 55.6 | 45.0 | | |
| CYP20A1_3'UTR_FP9 | TTTGTAAGAGCTTCAGGGAA | 20 | 54.8 | 40.0 | 209 | used as the "3'UTR primer" |
| CYP20A1_3'UTR_RP9 | CACCGAACTGTAAACCAATT | 20 | 54.7 | 40.0 | | |
| CYP20A1_3'UTR_FP10 | GCTTCAGGGAATAGGCTTTT | 20 | 56.3 | 45.0 | 177 | |
| CYP20A1_3'UTR_RP10 | TTGGGAGTAGAACTGGAAGA | 20 | 55.7 | 45.0 | | |
| CYP20A1_Exon5_FP | GGAAGTCATTCGCTTCCAGA | 20 | 64.3 | 50.0 | 277 (including exon6) & 196 (excluding exon 6) | for discrimating the transcript isoforms |
| CYP20A1_Exon8_RP | TATGCAACTGGCCAGAGAAA | 20 | 63.3 | 45.0 | | |
| MALAT1_FP | CTTCCTGTGGCAGGAGAGAC | 20 | 64.1 | 60.0 | | |
| MALAT1_RP | CGCTTGAGATTTGGGCTTTA | 20 | 64 | 45.0 | 217 | |
| GAPDH_FP | CGACCACTTTGTCAAGCTCA | 20 | 58.99 | 50.0 | | |
| GAPDH_RP | CTTCCTCTTGTGCTCTTGCTG | 21 | 59.77 | 52.4 | 138 | |
| Spike-in FP | CCTCTTGTATCTCACAGCTCAA | 22 | 54.1 | 45.5 | | zebrafish lncRNA durga IVT product |
| Spike-in RP | CTGGAGACAATAGAAGGCAAT | 21 | 52.2 | 42.9 | 141 | |
| ACTB_FP | GTCTTCCCCTCCATCGTG | 18 | 63.3 | 61.1 | | |
| ACTB_RP | GATGGGGTACTTCAGGGTGA | 20 | 63.8 | 55.0 | 126 | |
| 18S rRNA_FP | GGCCCTGTAATTGGAATGAGTC | 22 | 59.63 | 50 | | |
| 18S rRNA_RP | CCAAGATCCAACTACGAGCTT | 21 | 58.63 | 47.6 | 146 | |

#### Antibodies used in WB

| Sl.no. | Company | Catalogue no. | Antibody name | Diltuion | Incubation |
|---|---|---|---|---|---|
| 1 | abcam | 136094 | Rabbit polyclonal to *CYP20A1* | 1:500 | Overnight at 4°, without shaking |
| 2 | Santa cruz | sc32233 | Mouse monoclonal anti *GAPDH* | 1:2000 | RT for 2 hrs, with shaking |
| 3 | abcam | 176842 | Rabbit monoclonal anti H3 | 1:1000 | Overnight at 4°, with shaking |
| 4 | Santa cruz | sc 2004 | anti rabbit IgG-HRP conjugated rasied in goat | 1:10,000 | RT for 1 hour, with shaking |
| 5 | Santa cruz | sc 2005 | anti mouse IgG-HRP conjugated rasied in goat | 1:10,000 | RT for 1 hour, with shaking |

#### S3- Results

#### Details of 3’RACE amplicons

| Sl. No. | Band size (bp) | Primer | Seq. length (nt) | NCBI BLAST (hg38) | | | UCSC BLAT (hg38) |
|---|---|---|---|---|---|---|---|
| | | | | Top hit | Identity (%) | Query cover (%) | |
| 1 | ~1500 | FP10 | 546 | *CYP20A1* mRNA (NM_177538.2) | 99 | 34 | Mapped to *CYP20A1* 3'UTR with high score |
| 3 | >800 | FP10 | 181 | *CYP20A1* mRNA (NM_177538.2) | 98 | 100 | Mapped to *CYP20A1* 3'UTR with high score |
| 5 | >700 | FP10 | 656 | *CYP20A1* mRNA (NM_177538.2) | 99 | 28 | Mapped to *CYP20A1* 3'UTR with high score |
| 7 | >600 | FP10 | 592 | *CYP20A1* mRNA (NM_177538.2) | 98 | 32 | Mapped to Chr.6(-) with low score |
| 9 | ~500 | FP10 | 362 | *MFSD1* | 95 | 99 | Mapped to *MFSD1* exon 16 (3'UTR) with very high score |

* no significant similarity was found with inner primer for any of the bands

#### Sample-wise data generated in primary neurons RNA-seq experiment

| Sample Name | Read 1 | Read 2 | Total data (in GB) |
|---|---|---|---|
| Untreated Set 1 | 7.8 | 7.9 | 15.7 |
| Untreated Set 2 | 5.2 | 5.2 | 10.4 |
| Untreated Set 3 | 11.8 | 11.2 | 23 |
| HS treated + 1hr recovery Set 1 | 7.5 | 7.6 | 15.1 |
| HS treated + 1hr recovery Set 2 | 8.8 | 8.8 | 17.6 |
| HS treated + 1hr recovery Set 3 | 9.8 | 9.9 | 19.7 |
| Tat treated + with recovery Set 1 | 7.2 | 7.3 | 14.5 |
| Tat treated + with recovery Set 2 | 7.8 | 7.9 | 15.7 |
| Tat treated + with recovery Set 3 | 6.8 | 6.9 | 13.7 |

#### Details of small RNA sequencing data

| Sample | Total # of reads | % of 5'adapter +ve reads | % of 3'adapter +ve reads | # of reads post length filtering | # of reads post quality filtering | # of reads mapped |
|---|---|---|---|---|---|---|
| Pr. neuron Set 1 | 58507953 | 3.52 | 96.48 | 45873165 | 44297019 | 36653535 |
| Pr. neuron Set 2 | 65588157 | 4.07 | 95.93 | 52787771 | 51364925 | 40148127 |
| Pr. neuron Set 3 | 35409327 | 3.57 | 96.43 | 31974327 | 31426458 | 25742827 |
| MCF-7 Set 1 | 49174023 | 3.76 | 96.24 | 43824299 | 41480248 | 36657502 |
| MCF-7 Set 2 | 94370695 | 4.27 | 95.73 | 83124005 | 78450081 | 68786702 |
| MCF-7 Set 3 | 59179178 | 8.05 | 91.95 | 50024126 | 46832593 | 46832593 |
| MCF-7 Set 4 | 75156581 | 7.77 | 92.23 | 62389991 | 57937708 | 49243044 |
| MCF-7 Set 5 | 62465203 | 8.56 | 91.44 | 54622720 | 51028477 | 42745201 |
| MCF-7 Set 6 | 73825714 | 3.73 | 96.27 | 64923259 | 60749506 | 53921524 |

#### Details of nuclear cytosolic RT-qPCR

<table>
<tr><th rowspan="2">Experiment</th><th colspan="5">Lowest(#) Ct difference between +RT and -RT</th><th colspan="5">Ct value in NTC</th><th rowspan="2">Ct variation in Spike-in +RT(#)</th></tr>
<tr><th>3'UTR primer</th><th>5'UTR primer</th><th>MALAT1</th><th>GAPDH</th><th>Spike-in</th><th>3'UTR primer</th><th>5'UTR primer</th><th>MALAT1</th><th>GAPDH</th><th>Spike-in</th></tr>
<tr><td>1</td><td>ND</td><td>7.19</td><td>9.34</td><td>16.44</td><td>ND</td><td>ND</td><td>35.12</td><td>33.35*</td><td>39.5</td><td>35.39*</td><td>0.20</td></tr>
<tr><td>2</td><td>7.31</td><td>6.45</td><td>9.04</td><td>11.77</td><td>6.66</td><td>40.93*</td><td>33.99</td><td>33.3*</td><td>33.55</td><td>31.67*</td><td>0.06</td></tr>
<tr><td>3</td><td>2.87</td><td>7.40</td><td>8.93</td><td>13.03</td><td>4.88</td><td rowspan="2">32.03</td><td rowspan="2">32.56</td><td rowspan="2">33.49*</td><td rowspan="2">34.57</td><td rowspan="2">34.99*</td><td>0.58</td></tr>
<tr><td>4</td><td>3.81</td><td>6.71</td><td>9.56</td><td>14.13</td><td>4.85</td><td>1.09</td></tr>
</table>

Noise band was set at 1.00 in all experiments
### among the total, nuclear and cytosolic fractions
* Ct value detected in one technical replicate only
ND = Ct Not detected in -RT; NTC = no template control
RT-qPCR for both experiment 3 and 4 were put in the same plate, therefore single NTC value

#### Supplementary Table Legends

**Table S1**: 91 genes (out of 319 containing the cooccurrence of Alu exonization, A-to-I editing and cis-antisense at single exon level) were mapped onto the apoptosis network. Out of these, 68 clustered around three central hubs (termed ‘mito.’, ‘p53’ and ‘ubi’).

**Table S2:** RNA-seq reads from *CYP20A1* different transcript isoforms mapping to 15928 single nuclei from different regions of human cerebral cortex.

**Table S3:** Expression of *CYP20A1* in human specific Rosehip neurons.

**Table S4**: 46 out of the 169 miRNAs with MREs on *CYP20A1* 3'UTR were present in miRDB as FuncMir. 19 of them occur exclusively in one or more primate species; some of them are also stress responsive.

**Table S5**: List of 994 miRNAs with predicted MREs on *CYP20A1*_ALu-LT. Out of these 994, 140 miRNAs have ≥10 MREs.

**Table S6**: 380 genes that show concordant directionality of expression as *CYP20A1*_Alu-LT in heat shock and Tat response.

**Table S7**: Major biological pathways enriched in toppfun analysis of 380 genes using Benjamini-Hochberg corrected FDR (q- value 0.05).

**Supplementary Figure Legends**

**Figure S1**: Out of 38300 genes, 512 have their longest 3'UTR greater than 6kb but only 98 of these contain Alu exonization events.

**Figure S2**: Snapshot (1-60aa) of multiple sequence alignment of CYP20A1 proteins depicts high degree of sequence conservation across vertebrates. As expected, the primates show the highest similarity to the human protein, which reduces as we move from the mammals to other non-mammalian vertebrates. The similarity is least for zebrafish which is also the most evolutionarily distant vertebrate from humans. (* exact aa conserved, : replaced by aa containing similar functional group)

**Figure S3**: UCSC track representing DNA conservation analysis of 10kb upstream of 5'UTR and 10 kb downstream of 3'UTR region among 20 mammals.

**Figure S4**: Expression of *CYP20A1* isoform in MCF-7 cells was checked by RT-qPCR using two different primer pairs and normalized to the geometric mean of β-actin, *GAPDH* and 18S rRNA. The error bars represent the SD of four biological replicates. Average of three technical replicates was taken for each biological replicate.
In contrast to the low expression of our transcript of interest, CYP20A1 protein is highly expressed in MCF-7 cells (~50% of GAPDH level).

**Figure S5**: Transcript models built by Cufflinks based on RNA-seq data suggest that 3'UTRs longer than 9kb, occur for *CYP20A1*. TCONS_00101353 denotes the isoform studied by us.

**Figure S6**: >95% of the cells are immuno-positive for neuronal markers like MAP2, Tuj1 and NeuN, while no cells are stained with GFAP, a glial cell marker, ensuring it is a pure neuron culture.

**Figure S7**: **A.** ‘Kill-curve’ of different doses of HIV1-Tat on primary neurons was prepared using viability and cytotoxicity parameters (ApoToxGlo assay; Promega) after 24hrs of treatment.
**B.** Subsequently, the protocol was standardized for 6hrs treatment. ~40% cell death occurs in primary neurons upon 100ng/ml Tat treatment for 6hrs. The data represent the mean (±S.D.) of two independent experiments.

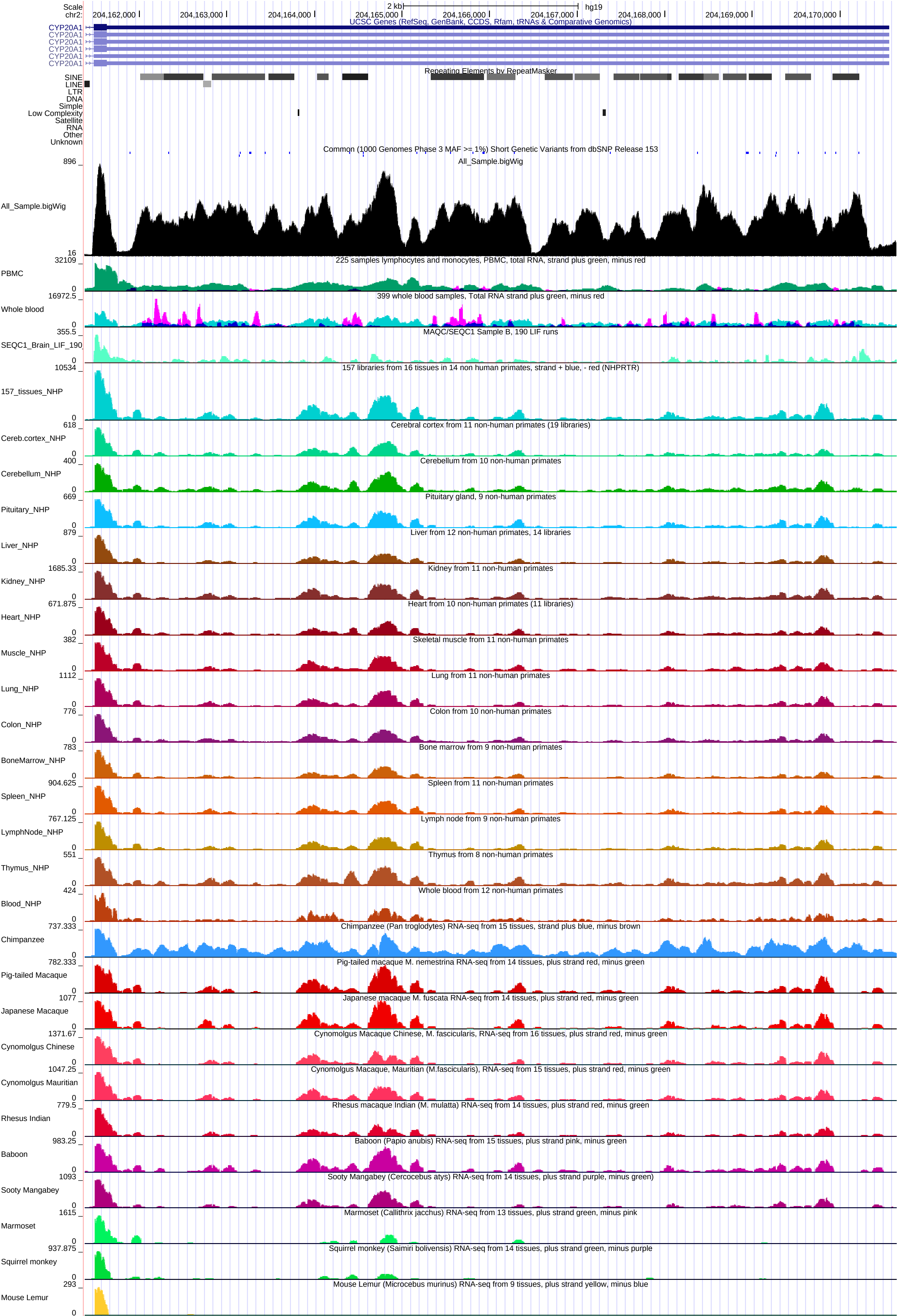

Scale
chr2:
2 kb
hg19
204,162,000
204,163,000
204,164,000
204,165,000
204,166,000
204,167,000
204,168,000
204,169,000
204,170,000
UCSC Genes (RefSeq, GenBank, CCDS, Rfam, tRNAs & Comparative Genomics)
CYP20A1
CYP20A1
CYP20A1
CYP20A1
CYP20A1
CYP20A1
Repeating Elements by RepeatMasker
SINE
LINE
LTR
DNA
Simple
Low Complexity
Satellite
RNA
Other
Unknown
Common (1000 Genomes Phase 3 MAF >= 1%) Short Genetic Variants from dbSNP Release 153
All_Sample.bigWig
896
All_Sample.bigWig
16
32109
PBMC
0
225 samples lymphocytes and monocytes, PBMC, total RNA, strand plus green, minus red
16972.5
Whole blood
0
399 whole blood samples, Total RNA strand plus green, minus red
355.5
SEQC1_Brain_LIF_190
0
MAQC/SEQC1 Sample B, 190 LIF runs
10534
157_tissues_NHP
0
157 libraries from 16 tissues in 14 non human primates, strand + blue, - red (NHPRTR)
618
Cereb.cortex_NHP
0
Cerebral cortex from 11 non-human primates (19 libraries)
400
Cerebellum_NHP
0
Cerebellum from 10 non-human primates
669
Pituitary_NHP
0
Pituitary gland, 9 non-human primates
879
Liver_NHP
0
Liver from 12 non-human primates, 14 libraries
1685.33
Kidney_NHP
0
Kidney from 11 non-human primates
671.875
Heart_NHP
0
Heart from 10 non-human primates (11 libraries)
382
Muscle_NHP
0
Skeletal muscle from 11 non-human primates
1112
Lung_NHP
0
Lung from 11 non-human primates
776
Colon_NHP
0
Colon from 10 non-human primates
783
BoneMarrow_NHP
0
Bone marrow from 9 non-human primates
904.625
Spleen_NHP
0
Spleen from 11 non-human primates
767.125
LymphNode_NHP
0
Lymph node from 9 non-human primates
551
Thymus_NHP
0
Thymus from 8 non-human primates
424
Blood_NHP
0
Whole blood from 12 non-human primates
737.333
Chimpanzee
0
Chimpanzee (Pan troglodytes) RNA-seq from 15 tissues, strand plus blue, minus brown
782.333
Pig-tailed Macaque
0
Pig-tailed macaque M. nemestrina RNA-seq from 14 tissues, plus strand red, minus green
1077
Japanese Macaque
0
Japanese macaque M. fuscata RNA-seq from 14 tissues, plus strand red, minus green
1371.67
Cynomolgus Chinese
0
Cynomolgus Macaque Chinese, M. fascicularis, RNA-seq from 16 tissues, plus strand red, minus green
1047.25
Cynomolgus Mauritian
0
Cynomolgus Macaque, Mauritian (M.fascicularis), RNA-seq from 15 tissues, plus strand red, minus green
779.5
Rhesus Indian
0
Rhesus macaque Indian (M. mulatta) RNA-seq from 14 tissues, plus strand red, minus green
983.25
Baboon
0
Baboon (Papio anubis) RNA-seq from 15 tissues, plus strand pink, minus green
1093
Sooty Mangabey
0
Sooty Mangabey (Cercocebus atys) RNA-seq from 14 tissues, plus strand purple, minus green)
1615
Marmoset
0
Marmoset (Callithrix jacchus) RNA-seq from 13 tissues, plus strand green, minus pink
937.875
Squirrel monkey
0
Squirrel monkey (Saimiri bolivensis) RNA-seq from 14 tissues, plus strand green, minus purple
293
Mouse Lemur
0
Mouse Lemur (Microcebus murinus) RNA-seq from 9 tissues, plus strand yellow, minus blue

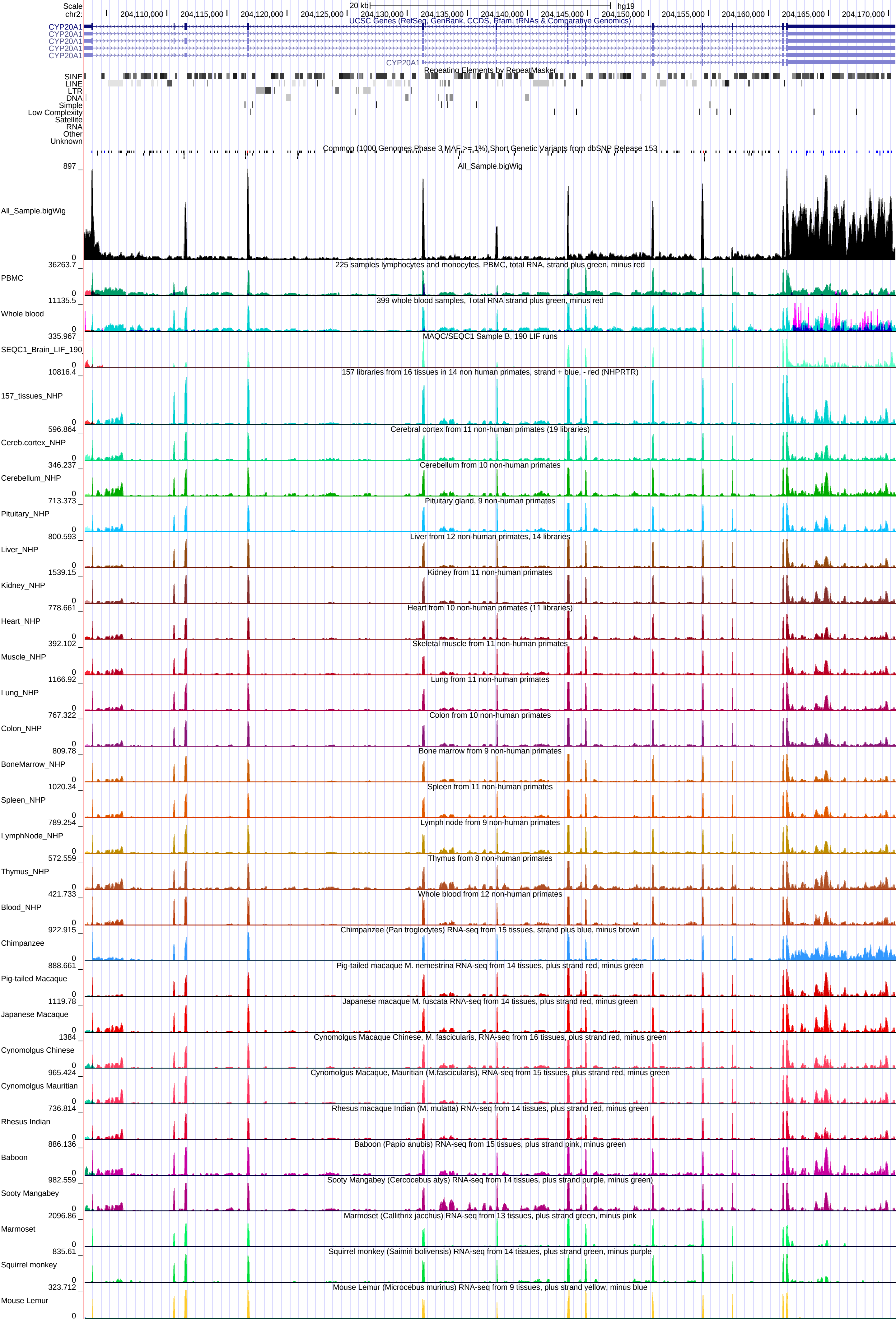
Scale
chr2:
20 kb
hg19
204,110,000
204,115,000
204,120,000
204,125,000
204,130,000
204,135,000
204,140,000
204,145,000
204,150,000
204,155,000
204,160,000
204,165,000
204,170,000
UCSC Genes (RefSeq, GenBank, CCDS, Rfam, tRNAs & Comparative Genomics)
CYP20A1
CYP20A1
CYP20A1
CYP20A1
CYP20A1
CYP20A1
Repeating Elements by RepeatMasker
SINE
LINE
LTR
DNA
Simple
Low Complexity
Satellite
RNA
Other
Unknown
Common (1000 Genomes Phase 3 MAF >= 1%) Short Genetic Variants from dbSNP Release 153
All_Sample.bigWig
897
0
All_Sample.bigWig
225 samples lymphocytes and monocytes, PBMC, total RNA, strand plus green, minus red
36263.7
PBMC
0
399 whole blood samples, Total RNA strand plus green, minus red
11135.5
Whole blood
0
MAQC/SEQC1 Sample B, 190 LIF runs
335.967
SEQC1_Brain_LIF_190
0
157 libraries from 16 tissues in 14 non human primates, strand + blue, - red (NHPRTR)
10816.4
157_tissues_NHP
0
Cerebral cortex from 11 non-human primates (19 libraries)
596.864
Cereb.cortex_NHP
0
Cerebellum from 10 non-human primates
346.237
Cerebellum_NHP
0
Pituitary gland, 9 non-human primates
713.373
Pituitary_NHP
0
Liver from 12 non-human primates, 14 libraries
800.593
Liver_NHP
0
Kidney from 11 non-human primates
1539.15
Kidney_NHP
0
Heart from 10 non-human primates (11 libraries)
778.661
Heart_NHP
0
Skeletal muscle from 11 non-human primates
392.102
Muscle_NHP
0
Lung from 11 non-human primates
1166.92
Lung_NHP
0
Colon from 10 non-human primates
767.322
Colon_NHP
0
Bone marrow from 9 non-human primates
809.78
BoneMarrow_NHP
0
Spleen from 11 non-human primates
1020.34
Spleen_NHP
0
Lymph node from 9 non-human primates
789.254
LymphNode_NHP
0
Thymus from 8 non-human primates
572.559
Thymus_NHP
0
Whole blood from 12 non-human primates
421.733
Blood_NHP
0
Chimpanzee (Pan troglodytes) RNA-seq from 15 tissues, strand plus blue, minus brown
922.915
Chimpanzee
0
Pig-tailed macaque M. nemestrina RNA-seq from 14 tissues, plus strand red, minus green
888.661
Pig-tailed Macaque
0
Japanese macaque M. fuscata RNA-seq from 14 tissues, plus strand red, minus green
1119.78
Japanese Macaque
0
Cynomolgus Macaque Chinese, M. fascicularis, RNA-seq from 16 tissues, plus strand red, minus green
1384
Cynomolgus Chinese
0
Cynomolgus Macaque, Mauritian (M.fascicularis), RNA-seq from 15 tissues, plus strand red, minus green
965.424
Cynomolgus Mauritian
0
Rhesus macaque Indian (M. mulatta) RNA-seq from 14 tissues, plus strand red, minus green
736.814
Rhesus Indian
0
Baboon (Papio anubis) RNA-seq from 15 tissues, plus strand pink, minus green
886.136
Baboon
0
Sooty Mangabey (Cercocebus atys) RNA-seq from 14 tissues, plus strand purple, minus green)
982.559
Sooty Mangabey
0
Marmoset (Callithrix jacchus) RNA-seq from 13 tissues, plus strand green, minus pink
2096.86
Marmoset
0
Squirrel monkey (Saimiri bolivensis) RNA-seq from 14 tissues, plus strand green, minus purple
835.61
Squirrel monkey
0
Mouse Lemur (Microcebus murinus) RNA-seq from 9 tissues, plus strand yellow, minus blue
323.712
Mouse Lemur
0
